# Inactivation of HIF-P4H-1 Stabilizes IKKα, Modulates Non-Canonical NF-κB Signaling, and Sensitizes Cancer Cells to Cell Death

**DOI:** 10.64898/2026.07.17.739273

**Authors:** Karim Ullah, Mizanur Rahman, Farhan Quadir, Valerio Izzi, Ulrich Bergmann, Jing Zhang, Qing Zhang, Rongxue Wu, Joni M. Mäki, Johanna Myllyharju

## Abstract

Hypoxia and NF-κB signaling are well-established drivers of cancer progression and treatment failure, yet the oxygen-dependent regulation of non-canonical NF-κB signaling remains poorly defined. Here, we identify hypoxia-inducible factor prolyl-4-hydroxylase-1 (HIF-P4H-1/EGLN2) as a key modulator of the non-canonical NF-κB pathway. Using integrated biochemical, genetic, proteomic, and transcriptomic analyses across human cell lines, mouse models, and clinical tumor samples, we demonstrate that HIF-P4H-1 directly interacts with and hydroxylates IKKα at proline 367, thereby promoting its ubiquitination and proteasomal degradation. Loss or inhibition of HIF-P4H-1 results in accumulation of IKKα, impaired NF-κB2/p100 processing to p52, destabilization of NF-κB-inducing kinase (NIK), and suppression of non-canonical NF-κB–dependent survival gene expression. Structural modeling and mutagenesis identify proline 367 hydroxylation as a critical determinant of IKKα turnover. Analysis of TCGA cohorts reveals an inverse correlation between HIF-P4H-1 and IKKα expression, with elevated HIF-P4H-1 associating with advanced tumor stage and reduced overall survival in clear cell renal cell carcinoma. Functionally, targeting HIF-P4H-1 sensitizes cancer cells to cell death and impairs proliferation, clonogenic growth, and migration. Together, our findings define a previously unrecognized oxygen-dependent mechanism regulating non-canonical NF-κB signaling through direct control of IKKα stability and nominate the HIF-P4H-1–IKKα axis as a potential therapeutic vulnerability in hypoxia-adapted malignancies.

## Introduction

Therapeutic resistance remains a major obstacle in cancer treatment [1, 2]. Lack of drug-induced apoptosis is a common mechanism by which cancer escapes from therapy [3]. The NF-κB family of transcription factors contributes to these processes by regulating genes involved in inflammation, proliferation, and anti-apoptotic responses [4, 5]. This family includes RelA, RelB, c-Rel, NF-κB1 (p105/p50), and NF-κB2 (p100/p52) which assemble into distinct transcriptional complexes to mediate two mechanistically separate pathways. The canonical NF-κB pathway mainly involves RelA:p50 dimers, responds rapidly to inflammatory stimuli, and activation relies on IKKβ-and IKKγ-dependent degradation of IκB proteins [6].

In contrast, the non-canonical NF-κB pathway is tightly regulated, and activation depends on NF-κB–inducing kinase (NIK) and IKKα-mediated phosphorylation of NFκB2/p100 [7], leading to NF-κB2/p100 processing to p52 and nuclear translocation of RelB:p52 heterodimers [7, 8]. In normal conditions, NF-κB2/p100 accumulates as the inactive precursor, and its processing occurs only upon stimulation of specific receptors such as lymphotoxin β, RANK, CD40, and BAFF [7, 9]. While NF-κB signaling is broadly implicated in therapy resistance, most studies have focused on canonical RelA/p50 signaling. Evidence supporting a role for the non-canonical NIK–IKKα–NF-κB2/p52 pathway remains limited and has been described mainly in specific tumor contexts such as multiple myeloma, where constitutive p100 processing promotes tumor cell survival and drug resistance [10, 11]. The IκB kinase (IKK) complex, consisting of the catalytic subunits IKKα and IKKβ and the regulatory subunit IKKγ, serves as a central hub for NF-κB activation [12]. Notably, in the non-canonical NF-κB pathway, IKKα functions independently of IKKβ and IKKγ, forming homodimers that phosphorylate the NF-κB2/p100 precursor and trigger its partial proteasomal processing into the active p52 subunit [13].

Although hypoxia and NF-κB signaling are well-established drivers of tumor progression and treatment failure, the role of non-canonical IKKα–NF-κB2/p52 signaling, particularly in hypoxic contexts, remains poorly understood. Prior studies demonstrate that hypoxia and prolyl hydroxylases regulate canonical NF-κB activity, suggesting potential crosstalk between oxygen sensing and NF-κB pathways [14–16].

Hypoxia-inducible factor (HIF) prolyl hydroxylases (HIF-P4Hs, also known as PHDs/EGLNs) are oxygen-sensing enzymes that catalyze proline hydroxylation of diverse substrates beyond HIF, including p53, FOXO3a, and Cep192 [17–19]. Through this hydroxylation-dependent regulation, HIF-P4Hs critically modulate the stability and activity of their target proteins. Hypoxic inhibition of HIF-P4Hs alters numerous signaling pathways, including NF-κB [20, 21]. Notably, inhibition of HIF-P4H-1 reduces cyclin D1 expression and suppresses proliferation in cancer cells [22, 23]. Yet, whether HIF-P4H-1 directly modulates the non-canonical NF-κB pathway has not been investigated. In this study, we identify a previously unrecognized link between hypoxia sensing and non-canonical-NF-κB signaling. Analysis of primary clear-cell renal cell carcinoma (ccRCC) patient tissues revealed decreased IKKα protein abundance in tumors, accompanied by enhanced p100 processing and increased cyclin-D1 expression. These findings suggest that HIF-P4H-1 may influence IKKα protein stability, thereby activating non-canonical NF-κB signaling within the hypoxic tumor microenvironment. To establish the generality of this regulatory mechanism, we investigated the HIF-P4H-1–IKKα axis across multiple complementary cell models and patient-derived tumor samples.

## Results

### HIF-P4H-1 expression is associated with reduced IKKα protein levels in ccRCC

To explore the relationship between HIF-P4H-1 (EGLN2) and IKKα (also known by its gene name CHUK) in cancer, we first analyzed protein expression in human clear cell renal cell carcinoma (ccRCC) tumor samples by western blotting. Compared with matched adjacent normal tissues, ccRCC tumors exhibited increased expression of the oxygen-sensing enzyme HIF-P4H-1 (Fig. 1a-b), accompanied by reduced IKKα protein levels (Fig. 1e-f). In addition, components associated with non-canonical NF-κB signaling, including altered p100/p52 processing, as well as the downstream target Cyclin D1, showed tumor-associated changes (Fig. 1c–d). Correlation analysis of paired ccRCC samples revealed a negative trend between HIF-P4H-1 and IKKα protein levels ( Pearson, r = -0.4611, p = 0.25) (Supplementary Fig. 1a). Notably, VHL-mutant ccRCC tumor samples did not show a consistent reduction in IKKα protein abundance (Fig. 1g). Von Hippel–Lindau (VHL) is an E3 ubiquitin ligase required for hydroxylation-dependent proteasomal degradation. Therefore, VHL-mutant tumors were analyzed separately, as impaired VHL activity may uncouple HIF-P4H-1–mediated hydroxylation from efficient ubiquitination and degradation of IKKα, potentially explaining the lack of consistent IKKα reduction in this subset. Together, these protein-level data indicate an inverse association between HIF-P4H-1 and IKKα protein levels in ccRCC and provide a clinical context for subsequent mechanistic analyses. To extend these observations, we analyzed publicly available data from The Cancer Genome Atlas (TCGA) for gene expression, tumor staging, and patient survival. Coexpression analysis in ccRCC (TCGA, Firehose Legacy) revealed a robust inverse correlation between EGLN2 (HIF-P4H-1) and CHUK (IKKα) mRNA abundance (Spearman r = –0.43, p = 5.49 × 10⁻²⁵; Pearson r = –0.49, p = 3.0 × 10⁻³³; Supplementary Fig. 1a). In addition, correlation analysis of HIF-P4H-1 (EGLN2) and IKKα protein levels in paired ccRCC tumor samples revealed a negative, but non-significant, association (Pearson r = −0.46, p = 0.25; Supplementary Fig. 1b). Survival analysis demonstrated that elevated EGLN2 expression was significantly associated with reduced overall survival in ccRCC patients (log-rank p = 1.7 × 10⁻⁶; Fig. 1h), with high EGLN2 conferring a two-fold greater hazard of mortality (HR = 2.0, p = 2.6 × 10⁻⁶; n = 400 per group). Conversely, lower CHUK expression correlated with poorer overall survival, whereas higher CHUK expression was associated with improved patient outcomes (log-rank p = 0.00037; HR(high) = 0.57, p = 0.00044; Fig. 1i). In contrast, EGLN2 and CHUK expression in breast cancer cohorts did not correlate with patient outcome (HR = 1.1, log-rank p = 0.71; n = 535 per group), underscoring potential tissue-specific roles for non-canonical NF-κB pathway components (Supplementary Fig. 1c-d). EGLN2 transcript levels increased significantly with advancing ccRCC tumor stage (ANOVA F = 35.7, p = 5.21 × 10⁻²⁹; Fig. 1j). Consistent with these findings, analysis of medulloblastoma cell models and human tumor specimens revealed higher HIF-P4H-1 expression accompanied by reduced IKKα protein levels in more aggressive disease states (Fig. 1k–l; Supplementary Fig. 1e). Collectively, these patient-derived datasets provide clinical and pathological context linking elevated EGLN2 expression with reduced IKKα abundance and tumor aggressiveness, while the mechanistic basis for this relationship is established in the cellular and biochemical analyses described below.

**Figure 1:**
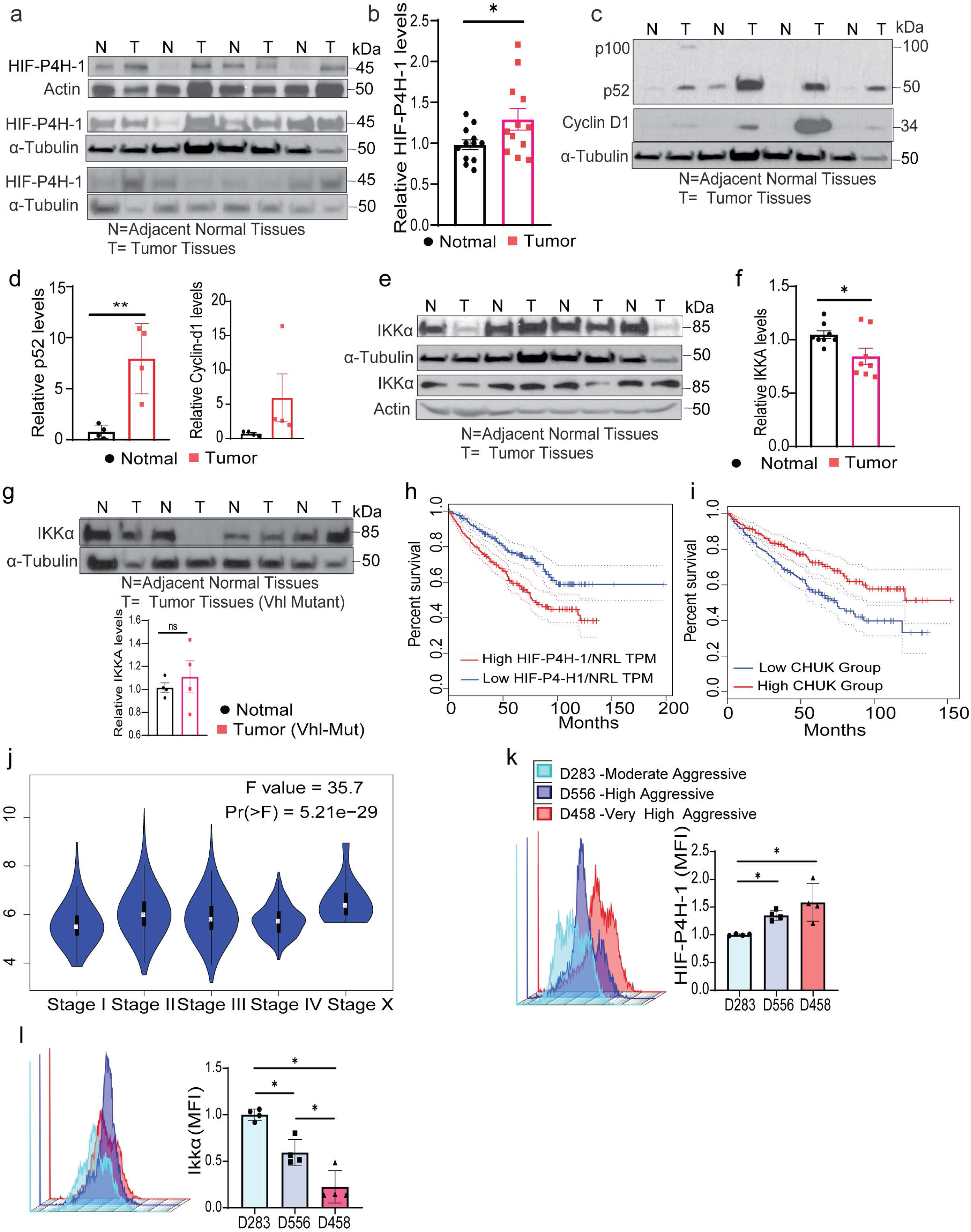
HIF-P4H-1 expression is associated with reduced IKKα protein levels in ccRCC. (a) Whole-tissue lysates from ccRCC tumors and matched adjacent normal tissues (n = 12) were analyzed by western blot using an HIF-P4H-1 antibody. (b) Densitometric quantification of HIF-P4H-1 protein abundance. (c, d) Western blot analysis and quantification of NF-κB2/p100–p52 and Cyclin D1 protein levels (n = 4). (e, f) Western blot analysis of IKKα protein abundance in the same ccRCC tumor–normal pairs, with corresponding quantification (n = 8). Figure 1a–g was generated from a total of three independent PVDF membranes. For panels derived from the same membrane, proteins were detected sequentially following stripping and re-probing; therefore, the same α-tubulin loading control is shown where applicable. The same control α-Tubulin blot is shown for panels c and e as the proteins were analyzed on the same membrane (g) IKKα protein levels in Vhl-mutant ccRCC tumors and matched normal tissues, with quantification shown below (n = 4). (h) Kaplan–Meier analysis of overall survival stratified by EGLN2 expression (n = 400 high; n = 400 low). High EGLN2 expression was associated with reduced overall survival (log-rank p = 1.7 × 10⁻⁶; HR = 2.0, p = 2.6 × 10⁻⁶). (i) Kaplan–Meier analysis of overall survival stratified by CHUK expression (n = 258 high; n = 257 low). High CHUK expression correlated with improved survival (log-rank p = 0.00037; HR = 0.57, p = 0.00044). Shaded areas indicate 95% confidence intervals. (j) Boxplot of EGLN2 expression across tumor stages (Stage I: n = 235; Stage II: n = 63; Stage III: n = 193; Stage IV: n = 66) showing significant stage-associated variation (ANOVA: F = 35.7, p = 5.21 × 10⁻²⁹). (k, l) Flow-cytometry analysis of HIF-P4H-1 and IKKα protein abundance in low-and high-aggressive medulloblastoma models (n = 4). For all comparisons, Student’s t-test was used. p < 0.05; *p < 0.01; **p < 0.001.

### Loss of HIF-P4H-1 stabilizes IKKα protein

To determine whether HIF-P4H-1 directly regulates IKKα protein stability, we next examined the effects of HIF-P4H-1 loss across multiple cell models. Consistent findings emerged following siRNA-mediated HIF-P4H-1 silencing in HEK293 cells, which resulted in increased IKKα protein levels (Fig. 2a). Similarly, mouse embryonic fibroblasts (MEFs) and macrophages from *Hif-p4h-1^-/-^* mice exhibited elevated IKKα protein levels compared to wild-type controls (Fig. 2b-c), whereas Cyclin D1 levels were reduced (Fig. 2b). Quantitative PCR analysis confirmed that the increased IKKα protein levels in *Hif-p4h-1^-/-^*MEFs and macrophages were not caused by elevated *Chuk* (*Ikkα)* mRNA transcription (Supplementary Fig. 2a). In vivo skin samples of TPA (12-O-tetradecanoylphorbol-13-acetate) treated *Hif-p4h-1^-/-^* mice showed a robust increase in IKKα protein compared to wild-type controls (Fig. 2d). To further evaluate the functional relationship between HIF-P4H-1 and IKKα, MDA-MB-231 breast cancer cells were treated with the pan-hydroxylase inhibitor FG4497, resulting in increased IKKα protein abundance (Fig. 2e). In contrast, IKKα protein levels were unchanged in Hif-p4h-3^-/-^ MEFs relative to wild-type controls (Fig. 2f). Likewise, genetic deletion of HIF-P4H-2 or HIF-P4H-3 in HEK293 cells did not alter IKKα protein abundance (Fig. 2g, h), supporting a selective role for HIF-P4H-1 in regulating IKKα stability.

**Figure 2:**
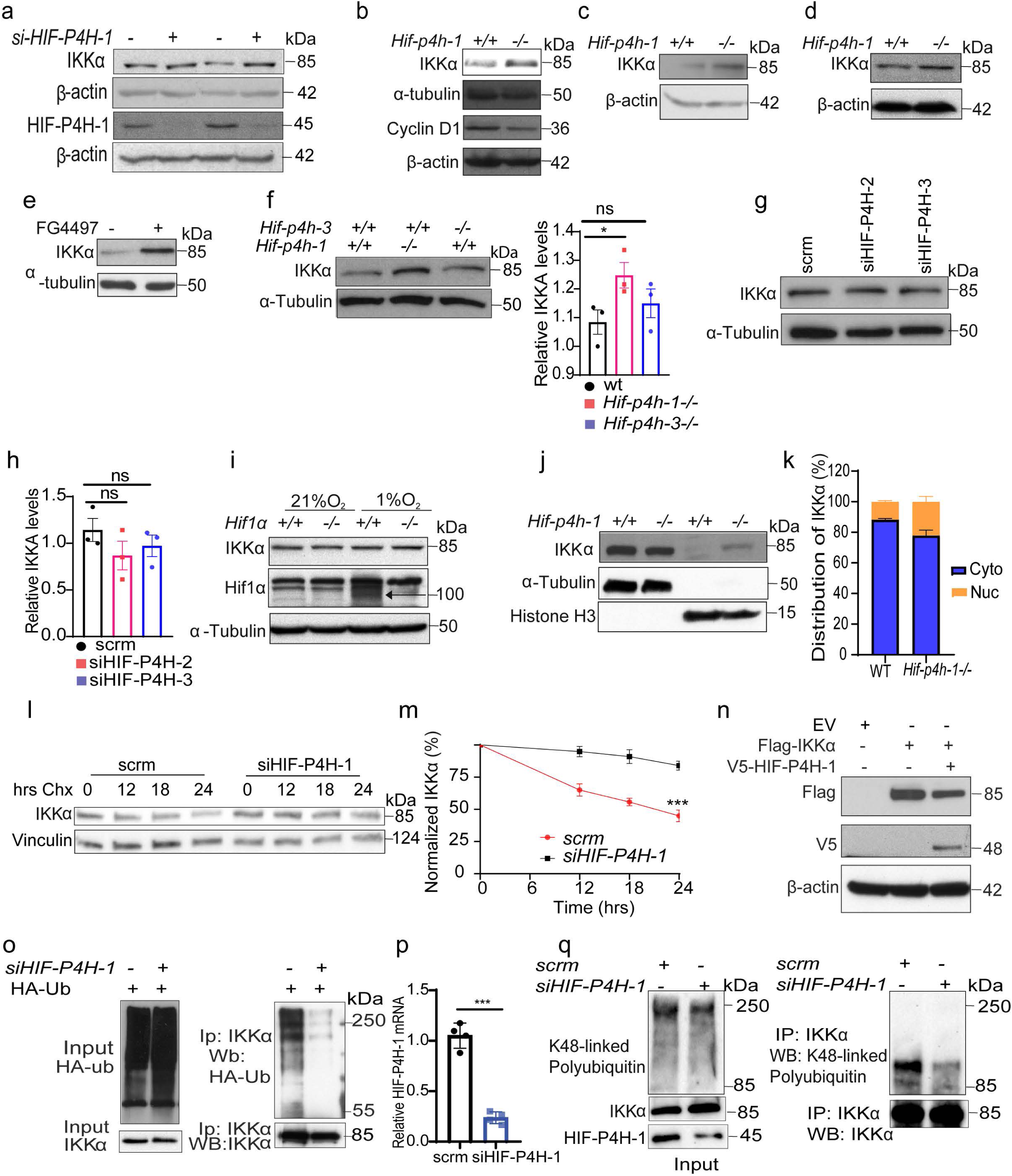
Loss of HIF-P4H-1 stabilizes IKKα. (a) siRNA-mediated silencing of HIF-P4H-1 in HEK293 cells increases endogenous IKKα protein levels, as assessed by western blot. (b–d) Western blot analysis of IKKα protein abundance in whole-cell lysates from Hif-p4h-1^-/-^and wild-type (WT) MEFs, primary macrophages, and skin tissues. (e) Pharmacological inhibition of HIF-P4H-1 with FG4497 (50 μM, 6 h) stabilizes IKKα protein levels in MDA-MB-231 cells (n = 4). (f) Western blot analysis and densitometric quantification of IKKα protein levels in WT, *Hif-p4h-1^-/-^*, and *Hif-p4h-3^-/-^*MEFs (n = 3). (g, h) Western blot analysis and quantification of IKKα protein levels following siRNA-mediated knockdown of HIF-P4H-2 or HIF-P4H-3 in HEK293 cells (n = 3). (i) Western blot analysis of IKKα protein levels in HIF1α^-/-^and WT HCT-116 cells cultured under normoxic (21% O₂) or hypoxic (1% O₂) conditions for 24 h (n = 3). (j, k) Western blot analysis of cytoplasmic and nuclear fractions from Hif-p4h-1^-/-^ and WT MEFs, and quantification of IKKα subcellular distribution (n = 3). (l, m) Cycloheximide chase assays (75 μM) reveal delayed IKKα degradation in HIF-P4H-1-depleted HCT-116 cells (n = 3). (n) Overexpression of V5-tagged HIF-P4H-1 reduces IKKα protein levels in MDA-MB-231 cells co-expressing FLAG-IKKα (n = 3). (o) HIF-P4H-1 promotes polyubiquitination of IKKα. HEK293 cells were transfected with HA-ubiquitin and treated with MG132, followed by siRNA-mediated knockdown of HIF-P4H-1. Endogenous IKKα was immunoprecipitated and ubiquitination was assessed by immunoblotting using anti-HA antibodies. (p) Quantitative RT–PCR analysis of HIF-P4H-1 mRNA levels confirming efficient knockdown under the same experimental conditions used for the ubiquitination assays. Data are shown as mean ± SD (n = 4). ***p < 0.001. (q) Representative immunoblot showing reduced K48-linked polyubiquitination of IKKα following siRNA-mediated knockdown of HIF-P4H-1 (n = 3 independent experiments).

To evaluate whether this regulation depends on canonical hypoxia signaling, HCT116 cells were treated with the pan-hydroxylase inhibitor DMOG followed by siRNA-mediated silencing of HIF1α; no significant changes in IKKα protein levels were observed (Supplementary Fig. 1d). Consistently, HIF1α^-/-^ and wild-type HCT116 cells exposed to hypoxia (1% O₂) showed comparable IKKα protein abundance (Fig. 2i), demonstrating that HIF-P4H-1–mediated regulation of IKKα is unaffected by HIF-1α expression. Subcellular fractionation further revealed increased accumulation of IKKα in the nuclear fraction of Hif-p4h-^1-/-^ MEFs compared with wild-type cells, without a corresponding increase in the cytoplasmic fraction (Fig. 2j, k). Cycloheximide chase assays demonstrated a prolonged half-life of IKKα following HIF-P4H-1 knockdown, confirming enhanced protein stability (Fig. 2l, m). Conversely, co-transfection of HIF-P4H-1 with IKKα in MDA-MB-231 cells resulted in a pronounced reduction in IKKα protein abundance compared with empty vector controls (Fig. 2n), consistent with accelerated protein turnover. Importantly, siRNA-mediated depletion of HIF-P4H-1 in cells co-expressing HA-tagged ubiquitin markedly reduced IKKα polyubiquitination relative to scrambled siRNA controls (Fig. 2o, and Supplementary Fig. 2c). To confirm knockdown efficiency, we quantified HIF-P4H-1 mRNA levels by qPCR. siRNA-mediated depletion of HIF-P4H-1 resulted in an 80–90% reduction in HIF-P4H-1 transcript abundance under the same conditions used for protein and functional assays (Fig. 2p). Similarly, K48-linked ubiquitination assays demonstrated a robust decrease in proteasome-targeting polyubiquitination of IKKα upon HIF-P4H-1 loss (Fig. 2q). Collectively, these data provide direct mechanistic evidence that HIF-P4H-1 selectively promotes ubiquitin-dependent degradation of IKKα, thereby regulating IKKα protein stability and turnover independently of HIF-1α under these experimental conditions.

### Structural analysis of IKKα interactions with HIF-P4H-1/2 and proximity of Pro367

To assess the potential for direct modification of IKKα by HIF-P4H-1, we employed AlphaFold2-Multimer to model the interaction between an IKKα peptide encompassing residues 356–377, including Pro367, and the catalytic domains of HIF-P4H-2 (Fig. 3a) and HIF-P4H-1 (Fig. 3b). The resulting in silico models revealed that Pro367 is located within a structurally flexible region of IKKα and is positioned at a substantial distance from the Fe(II)-coordinating His–Asp–His catalytic triad—12.2 Å for HIF-P4H-2 and 11.5 Å for HIF-P4H-1 (Fig. 3c, d). These distances exceed the typical catalytic range (<5–7 Å) required for efficient prolyl hydroxylation. Consistent with a weak or transient interaction, interface confidence scores were low (iPTM 0.32–0.41), and mean pLDDT values for the IKKα peptide were below 50, suggesting poor structural confidence and a lack of a catalytically optimized binding orientation. Substitution of Pro367 with alanine further disrupted peptide positioning and interface geometry (Fig. 3e, f), supporting a critical role for Pro367 in IKKα engagement with HIF-P4Hs. Collectively, these computational analyses indicate that while IKKα can transiently associate with HIF-P4H-1 and HIF-P4H-2, the modeled configurations do not support efficient hydroxylation of Pro367 in isolation. Productive modification may therefore require additional factors, such as cofactor binding, conformational rearrangements, or higher-order protein complexes, not captured in the static AlphaFold models.

**Figure 3:**
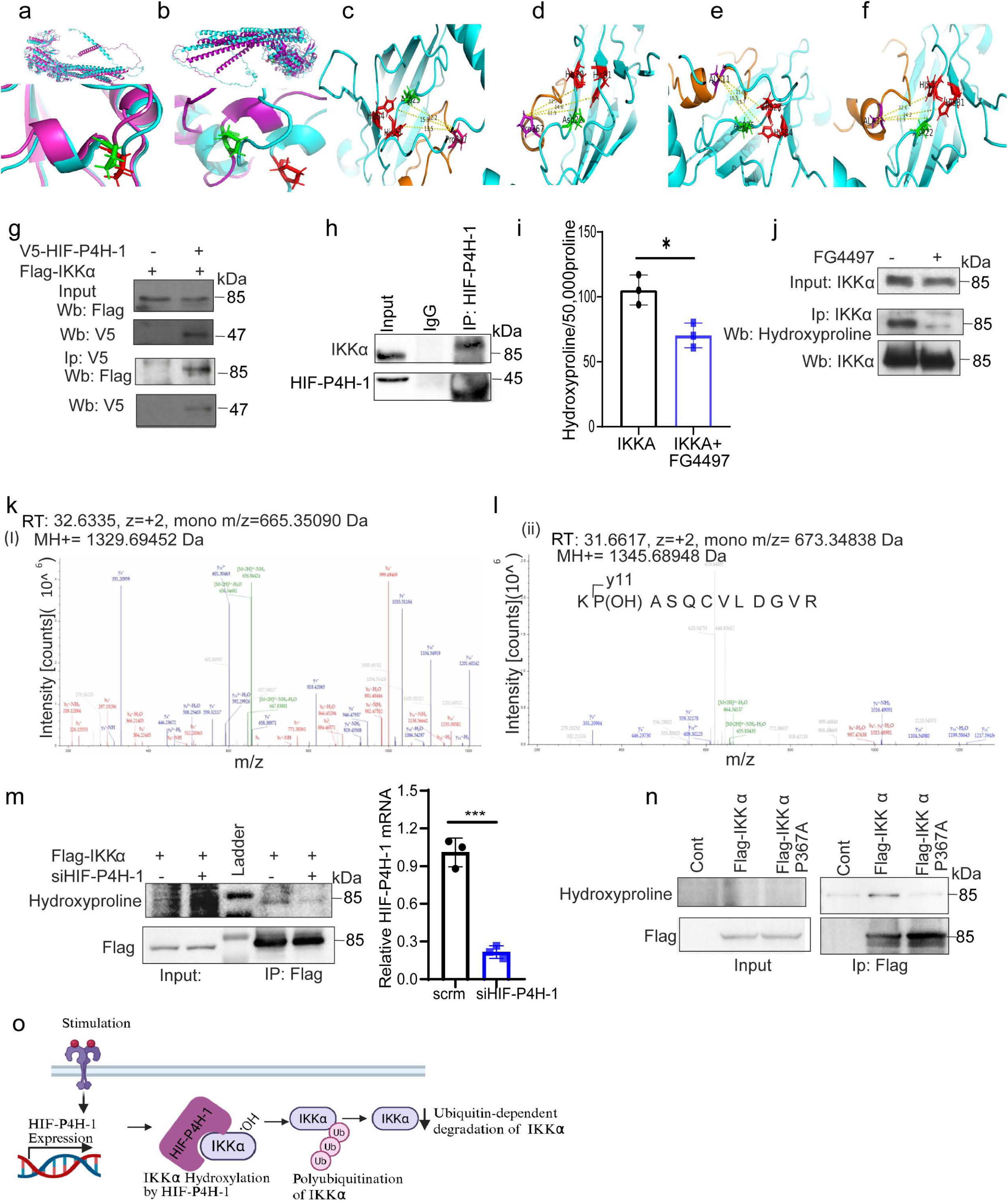
Structural and biochemical evidence for HIF-P4H-1–mediated hydroxylation of IKKα at Pro367. (a–f) Structural modeling of IKKα interactions with HIF-P4H-1 and HIF-P4H-2 and spatial positioning of Pro367. Superimposed AlphaFold2-predicted structures of full-length IKKα as a monomer (cyan) and in complex with HIF-P4H-2 (a) or HIF-P4H-1 (b) (purple); lower panels show zoomed views of Pro367, with the proline ring highlighted in green for the complexed form and red for the monomer. (c, d) Relative positioning of Pro367 (purple) within a 21-mer IKKα peptide (orange) in relation to the His–Asp–His catalytic triad (red–green–red) of HIF-P4H-2 (c) or HIF-P4H-1 (d) (cyan). Distances between Pro367 and the catalytic triad range from 11–16 Å, indicating a recognition-competent but catalytically suboptimal configuration. (e, f) Distance measurements for an alanine-substituted IKKα peptide (Pro367A) complexed with HIF-P4H-2 (e) or HIF-P4H-1 (f), showing no significant change in spatial arrangement or predicted catalytic competence. (g) HEK293 cells co-transfected with FLAG–IKKα and V5–HIF-P4H-1 were subjected to V5 immunoprecipitation, and interaction with IKKα was detected by anti-FLAG immunoblotting. (h) Endogenous HIF-P4H-1 was immunoprecipitated, followed by immunoblotting for IKKα to confirm endogenous interaction. (i) In vivo hydroxylation of IKKα assessed using [2,3,4,5 - ³H]-proline–labeled FLAG–IKKα expressed in MDA-MB-231 cells treated with or without FG4497 (50 μM); hydroxylation was quantified as 4-hydroxy[³H]proline per 50,000 proline residues (mean ± SD, n = 3). (j) MDA-MB-231 cells expressing FLAG–IKKα were treated with MG132 with or without FG4497, followed by FLAG immunoprecipitation and immunoblotting for IKKα and hydroxyproline. (k, l) LC–MS/MS identification of hydroxylation at Pro367 in FLAG–IKKα immunoprecipitated from MG132-treated MDA-MB-231 cells; MS/MS spectra of the peptide KPASQCVLDGVR (residues 366–377) show full y-ion coverage, with y11 confirming oxidation at Pro367. (m) siRNA-mediated silencing of HIF-P4H-1 in HEK293 cells reduces IKKα hydroxylation following FLAG–IKKα immunoprecipitation. (n) Overexpression of FLAG-tagged wild-type or Pro367A-mutant IKKα in HEK293 cells demonstrates loss of hydroxylation in the Pro367A variant following MG132 treatment. (o) Proposed model illustrating HIF-P4H-1 interaction with IKKα, Pro367 hydroxylation, and consequent ubiquitin-mediated degradation.

To assess the physical interaction between HIF prolyl hydroxylases and IKKα, we performed reciprocal co-immunoprecipitation assays in HEK293 cells co-transfected with Flag-tagged IKKα and V5-tagged HIF-P4H isoforms (HIF-P4H-1, -2, and 3). IKKα interacted with both HIF-P4H-1 and HIF-P4H-2, with the strongest association observed for HIF-P4H-1 (Fig. 3g, h; Supplementary Fig. 3a), identifying HIF-P4H-1 as a predominant binding partner of IKKα. In contrast, no interaction was detected between IKKα and HIF-P4H-3 (Supplementary Fig. 3a). Consistently, endogenous immunoprecipitation of IKKα from HCT116 cell lysates confirmed its interaction with HIF-P4H-1 (Fig. 3h). Together, these biochemical assays establish a selective physical association between HIF-P4H-1 and IKKα, supporting a direct role for HIF-P4H-1 in the regulation of IKKα protein stability.

Previous studies have shown that HIF-P4Hs can hydroxylate non-HIF substrates within consensus motifs such as -Leu-X-X-Leu-Ala-Pro-(Moser et al. 2013, Scholz & Taylor 2013, Ullah et al. 2017, Xie et al. 2012, Zheng et al. 2014b). Sequence analysis confirmed that both human and mouse IKKα harbor a conserved LXXLAP motif (LQYLAP, residues 185–190; Supplementary Fig. 3b), suggesting that IKKα may represent a potential non-HIF substrate of HIF-P4Hs. Based on these findings, we hypothesized that HIF-P4H-1 directly catalyzes the hydroxylation of IKKα. To test this, we performed in vivo hydroxyproline assays: MDA-MB-231 cells were transfected with Flag-tagged IKKα, labeled with L-[2,3,4,5-³H] proline, and treated with or without FG4497 and MG132. Immunoprecipitation followed by [³H]-4-hydroxyproline quantification showed significant incorporation of hydroxyproline into IKKα, which was markedly reduced by FG4497 treatment (Fig. 3i). Consistently, immunoblotting with an anti-hydroxyproline antibody confirmed robust IKKα hydroxylation, diminished upon HIF-P4H inhibition (Fig. 3j).

### HIF-P4H-1 Targets IKKα for Hydroxylation at Pro367

To identify site-specific hydroxylation of IKKα, Flag-tagged IKKα immunoprecipitates were resolved by SDS–PAGE, subjected to in-gel tryptic digestion, and analyzed by LC–MS/MS. This analysis achieved approximately 56% sequence coverage and identified a peptide spanning residues 366–377 (KPASQCVLDGVR) containing Pro367 in both hydroxylated and non-hydroxylated forms (Fig. 3k, l). Notably, exposure to hypoxic conditions resulted in an approximately 70% reduction in Pro367 hydroxylation (Supplementary Fig. 3c, d). Consistently, siRNA-mediated depletion of HIF-P4H-1 significantly reduced IKKα hydroxylation at this site (Fig. 3m).

To confirm site specificity, Flag-tagged wild-type IKKα or a Pro367Ala mutant was co-expressed in HEK293 cells. Hydroxylation was readily detected in the wild-type protein but was completely abolished in the Pro367Ala mutant (Fig. 3n), establishing Pro367 as the principal hydroxylation site targeted by HIF-P4H-1.

Together, these findings identify IKKα as a bona fide non-HIF substrate of HIF-P4H-1 and demonstrate that HIF-P4H-1–mediated hydroxylation at Pro367 represents a key post-translational modification associated with enhanced IKKα ubiquitination and proteasomal turnover, as summarized in the proposed model (Fig. 3o).

### HIF-P4H-1 Regulates NF-κB2/p100 Expression and Activity

To evaluate the clinical relevance of HIF-P4H-1 and NF-κB2 dysregulation in cancer, we analyzed transcriptomic data from The Cancer Genome Atlas breast cancer cohort (TCGA-BRCA; tumors n = 1085, normal tissues n = 112). Boxplot analyses revealed significant differential expression of both NFKB2 and HIF-P4H-1 (EGLN2) between tumor and normal tissues (Fig. 4a, b). Comparison of overall survival between high and low NFKB2 expression groups showed no statistically significant difference (log-rank p = 0.14; HR = 1.3, p = 0.14; Fig. 4c), whereas patients harboring EGLN2 alterations exhibited significantly worse survival outcomes (HR = 1.318, 95% CI: 1.007–1.724; log-rank p = 0.0161; Supplementary Fig. 4). In contrast, *NFKB2* alterations showed a non-significant trend toward improved survival (HR = 0.769, 95% CI: 0.587–1.008). Given that NF-κB2/p100 signaling is regulated predominantly at the post-translational level, alterations in *NFKB2* expression alone may not directly reflect non-canonical NF-κB pathway activity. To examine the functional impact of HIF-P4H-1 loss on NF-κB2 signaling, we measured p100/p52 protein levels in macrophages derived from *Hif-p4h-1^-/-^*mice. Compared with wild-type controls, *Hif-p4h-1^-/-^* macrophages exhibited a modest, non-significant reduction in p100 protein levels (Fig. 4d), without changes in Nfκb2 mRNA abundance (Supplementary Fig. 5a), consistent with post-transcriptional regulation. Despite limited changes in total p100 protein, expression of p52 target genes, including *Skp2* and *Bcl-xL*, was markedly reduced in *Hif-p4h-1^-/-^* macrophages (Fig. 4e). Similarly, *Bcl-xL* and *Vcam-1* were downregulated in *Hif-p4h-1^-/-^* MEFs (Supplementary Fig. 5b). Consistent results were observed in human cancer cells. siRNA-mediated depletion of HIF-P4H-1 in MDA-MB-231 cells reduced p100 protein abundance and significantly downregulated multiple p52-responsive genes, including SKP2, CCND1, BCL-XL, and CIAP-1, whereas VCAM-1 did not reach statistical significance (Fig. 4f, g; Supplementary Fig. 5c). Approximately 75% knockdown efficiency was confirmed by qPCR (Supplementary Fig. 5d). HIF-P4H-1 depletion also attenuated NF-κB transcriptional activity, as shown by reduced NF-κB luciferase reporter activity (Supplementary Fig. 5e). To assess upstream regulation of non-canonical NF-κB signaling, luciferase reporter assays were performed following NIK overexpression. NIK-driven NF-κB activation was markedly inhibited by *HIF-P4H-1* knockdown (Fig. 4h) and was similarly suppressed by treatment with the pan-hydroxylase inhibitor FG4497 (Fig. 4i). To explore potential feedback involving IKKα, we co-expressed IKKα and HIF-P4H-1 in MDA-MB-231 cells. While IKKα overexpression alone reduced NF-κB reporter activity, this effect was further enhanced by FG4497 treatment; conversely, HIF-P4H-1 overexpression increased NF-κB transcriptional activity (Fig. 4j). Collectively, these data support a role for HIF-P4H-1 in regulating non-canonical NF-κB signaling through effects on IKKα stability and NF-κB2/p100 processing in cancer cells.

**Figure 4:**
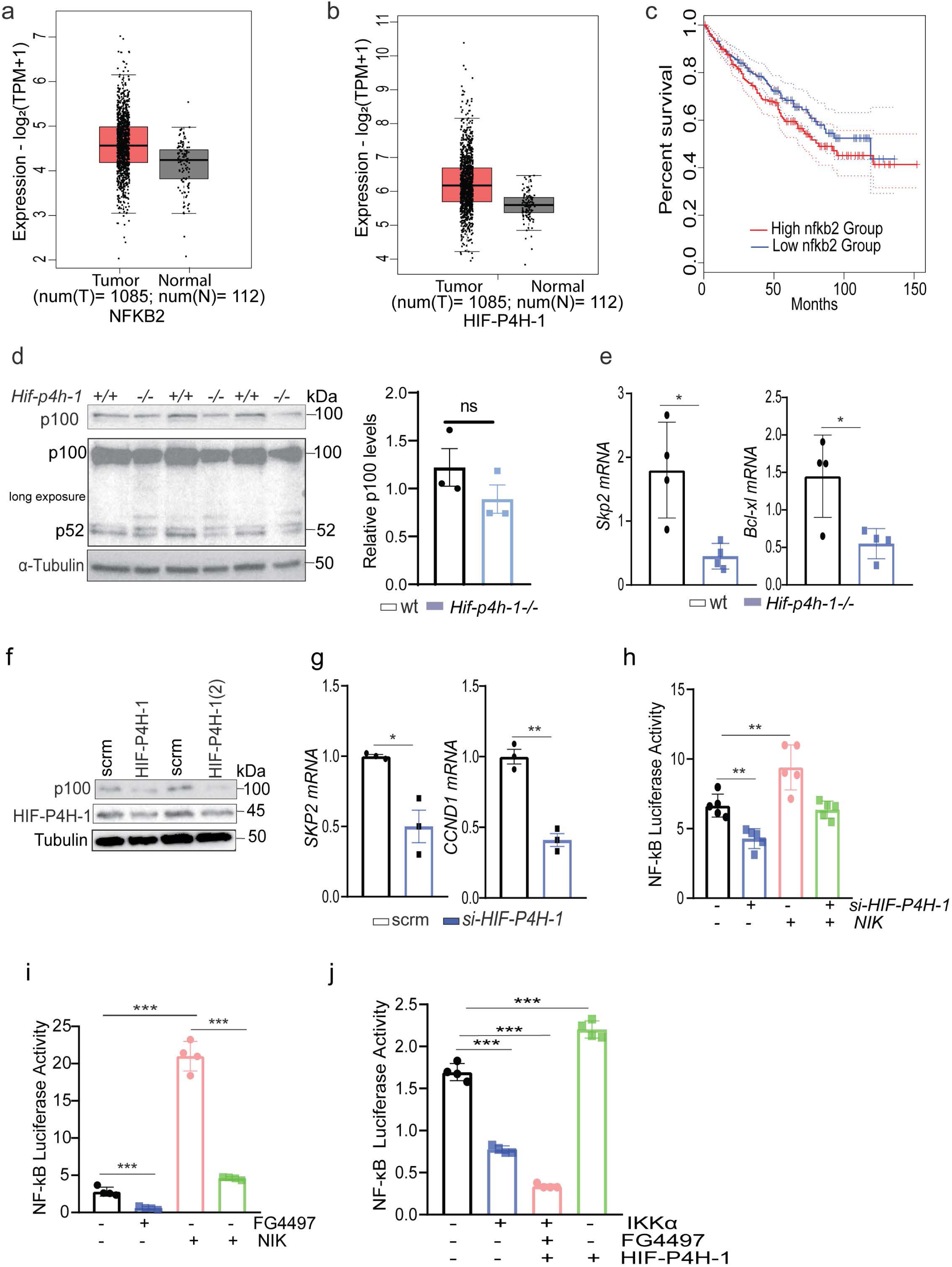
HIF-P4H-1 Regulates NF-κB2/p100 Expression and Activity. (a, b) Boxplot depicting NF-κB2 and HIF-P4H-1 expression levels in the specified study cohorts. The plot highlights indicated NF-κB2 and HIF-P4H-1 distribution (median, interquartile range, and outliers) across comparison groups. (c) Kaplan-Meier curves comparing overall survival between patients with high (n = 258) and low (n = 258) NF-κB2 expression. (d) Whole cell lysates from Hif-p4h-1-/-and WT macrophages were analyzed by WB for p100. (e) qPCR analysis of mRNA levels for Skp2 and Bcl-xl in Hif-p4h-1-/-and WT macrophages. (f) p100 and HIF-P4H-1 protein levels were analyzed from HIF-P4H-1 siRNA transfected MDA-MB-231 cells. (g) SKP2 and CYCLIN-D-1 mRNA expressions were analyzed by qPCR from HIF-P4H-1 siRNA-transfected cells. (h) NFκB luciferase reporter activity in HIF-P4H-1 knockdown and NIK-transfected MDA-MB-231 cells. (i) NFκB luciferase reporter activity in MDA-MB-231 cells. Cells were transfected with NIK and treated with or without FG4497 for 6h. (j) MDA-MB-231 cells were co-transfected with IKKα and HIF-P4H-1 and treated with or without FG4497 for 6h, and NFκB luciferase activity was measured. Data are presented as mean ± SD (n = 3-4). *p<0.05, **p<0.01 and ***p<0.001

### HIF-P4H-1 maintains IKKα-NIK homeostasis to enable inducible p100 processing

To investigate the role of HIF-P4H-1 in inducible NF-κB2/p100 processing, we first assessed IKKα protein abundance in MDA-MB-231 cells treated with the pan-hydroxylase inhibitor FG4497 and/or subjected to NIK overexpression. Inhibition of HIF-P4H-1 resulted in accumulation of IKKα protein, even in the presence of NIK (Fig. 5a), whereas NIK overexpression alone reduced IKKα abundance (Fig. 5a). Moreover, FG4497 treatment attenuated NIK-induced p100 processing, as evidenced by reduced p52 protein levels (Fig. 5b) and a modest but statistically significant attenuation of the NIK-induced increase in the p52 target gene Cyclin D1 (CCND1) mRNA (Fig. 5c). These data indicate that accumulation of IKKα following HIF-P4H-1 inhibition interferes with NIK-dependent p100-to-p52 processing.

**Figure 5:**
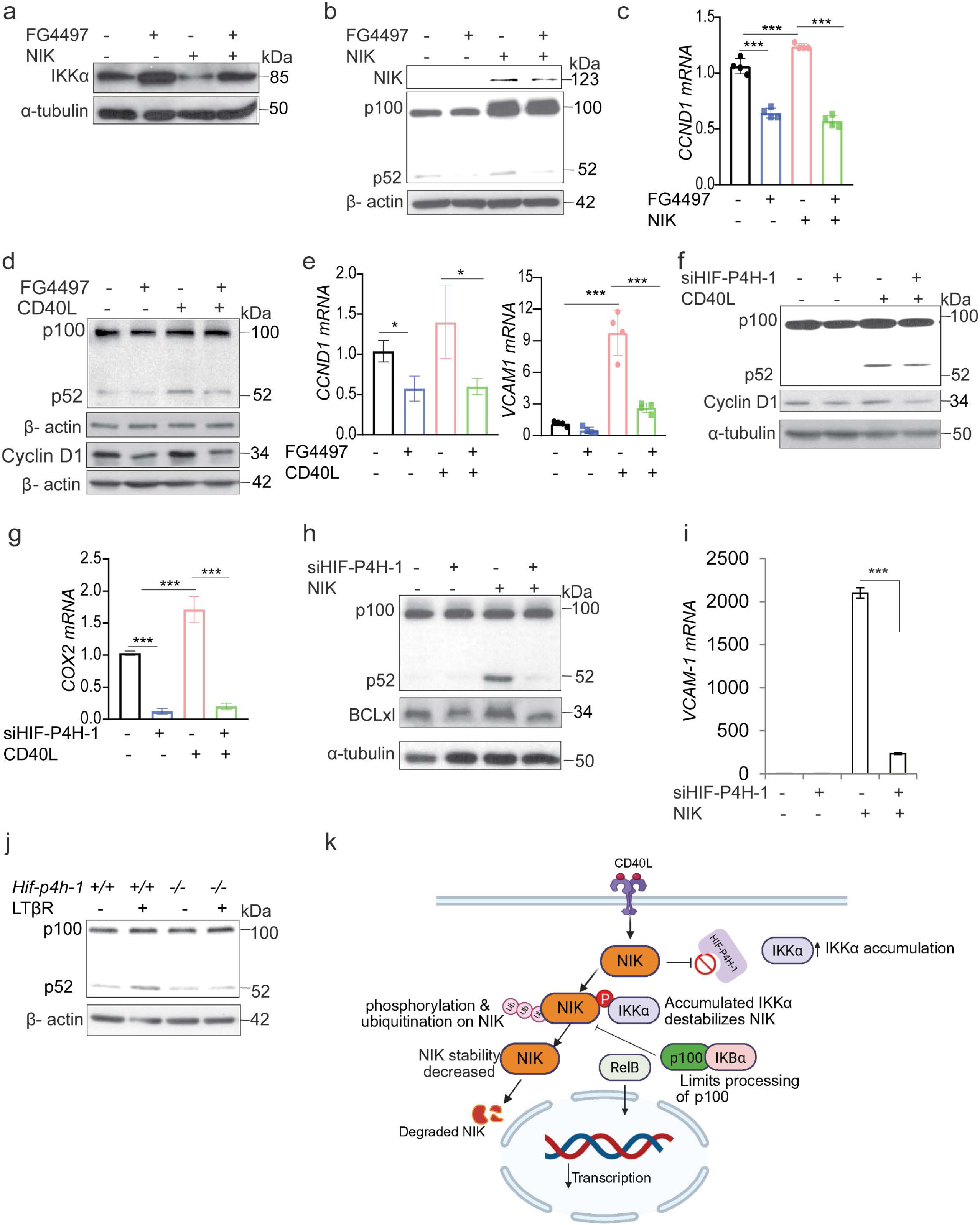
HIF-P4H-1 maintains IKKα–NIK homeostasis to enable inducible NF-κB2/p100 processing.(a–c) MDA-MB-231 cells were transfected with the indicated plasmids and treated with FG4497 (50 µM, 6 h). IKKα and NIK protein levels (a), NF-κB2/p100–p52 processing (b), and CCND1 (Cyclin D1) mRNA expression (c) were analyzed by western blot and qRT–PCR, respectively. (d, e) FG4497 treatment suppresses CD40L-induced p100 processing to p52: MDA-MB-231 cells were treated with FG4497 and stimulated overnight with soluble recombinant human CD40 ligand (hCD40L), followed by western blot analysis of indicated proteins (d) and qPCR analysis of CCND1 (Cyclin D1) and VCAM1 mRNA expression (e). (f, g) MDA-MB-231 cells transfected with HIF-P4H-1 siRNA were stimulated overnight with hCD40L, and p100/p52 and Cyclin D1 protein levels were analyzed by western blot, with COX2 mRNA expression quantified by qPCR (g). (h) HIF-P4H-1–depleted MDA-MB-231 cells were transfected with NIK, followed by western blot analysis of p100/p52 and qPCR analysis of VCAM1 mRNA expression (i). (j) *Hif-p4h-1^-/-^* MEFs were stimulated with LTβR agonist and analyzed by western blot for p100/p52 processing. (j) Schematic model illustrating that HIF-P4H-1 inactivation disrupts IKKα–NIK balance, resulting in impaired p100 processing to p52 and reduced expression of p52-dependent target genes.

Consistent with these findings, activation of non-canonical NF-κB signaling by CD40L stimulation also resulted in reduced p52 and Cyclin D1 protein levels in FG4497-treated cells (Fig. 5d). Analysis of p52-responsive gene expression, including CYCLIN D1 and VCAM-1, further confirmed impaired transcriptional induction in HIF-P4H-1–deficient cells following CD40L stimulation (Fig. 5e). Similarly, siRNA-mediated silencing of HIF-P4H-1 prevented CD40L-induced p100 processing to p52, as indicated by diminished p52 and Cyclin D1 protein levels (Fig. 5f). Reduced expression of the p52 target gene Cox2 further supported impaired p100-to-p52 processing under these conditions (Fig. 5g). Consistently, analysis of NF-κB2/p100 and BCL-xL protein levels, as well as VCAM-1 mRNA expression in HIF-P4H-1 silenced and NIK-overexpressing cells, demonstrated attenuated p100 processing to p52 and reduced downstream gene expression (Fig. 5h, i). Likewise, stimulation of Hif-p4h-1^-/-^ MEFs with LTβR ligand resulted in diminished p100-to-p52 processing compared with wild-type MEFs, providing additional biochemical evidence of impaired non-canonical NF-κB activation in the absence of HIF-P4H-1 (Fig. 5j). Notably, co-expression of IKKα and NIK resulted in a significant reduction in VCAM-1 mRNA levels (Supplementary Fig. 6a). Similarly, IKKα overexpression significantly attenuated CD40L-induced VCAM-1 expression (Supplementary Fig. 6b). In contrast, HIF-P4H-1 overexpression modestly enhanced CD40L-induced p100 processing to p52 (Supplementary Fig. 6c). Altogether, these data show that HIF-P4H-1 inhibition limits inducible p100-to-p52 processing and downstream non-canonical NF-κB transcription, as summarized in the proposed model (Fig. 5k).

### Inhibition of HIF-P4H-1 sensitizes cells to death by attenuating NF-κB2/p100 processing

Our data demonstrate that HIF-P4H-1 suppresses IKKα, thereby regulating NF-κB2/p100 processing and downstream transcription of pro-survival genes such as Cyclin D1 and Bcl-xL. To assess how HIF-P4H-1 influences cell proliferation and survival, transcriptomic analysis of Hif-p4h-1^-/-^ mouse embryonic fibroblasts (MEFs) revealed marked downregulation of proliferation-associated gene signatures under both normoxic and hypoxic conditions (Supplementary Table 1). Consistently, genetic deletion of HIF-P4H-1 in MEFs, siRNA-mediated knockdown, and pharmacological inhibition using FG4497 in MDA-MB-231 cells significantly reduced cellular proliferation rates (Fig. 6a–c). To determine whether HIF-P4H-1 modulates CD40L-induced non-canonical NF-κB signaling and downstream proliferative responses, proliferation assays were performed following HIF-P4H-1 inhibition. CD40L stimulation promoted proliferation of MDA-MB-231 cells, whereas FG4497 treatment markedly attenuated this effect (Fig. 6d). Likewise, overexpression of IKKα suppressed cell proliferation relative to controls (Fig. 6e), consistent with impaired NF-κB2/p100 signaling. To independently validate the effects of HIF-P4H-1 and IKKα on proliferation in a clinically distinct cancer context, Ki67 expression was analyzed by flow cytometry in medulloblastoma cells. Depletion of HIF-P4H-1 markedly reduced cancer cell proliferation. In contrast, IKKα silencing resulted in significantly higher Ki67 expression compared with HIF-P4H-1 depletion, although Ki67 levels remained lower than in scrambled siRNA controls (Supplementary Fig. 8c).

**Figure 6:**
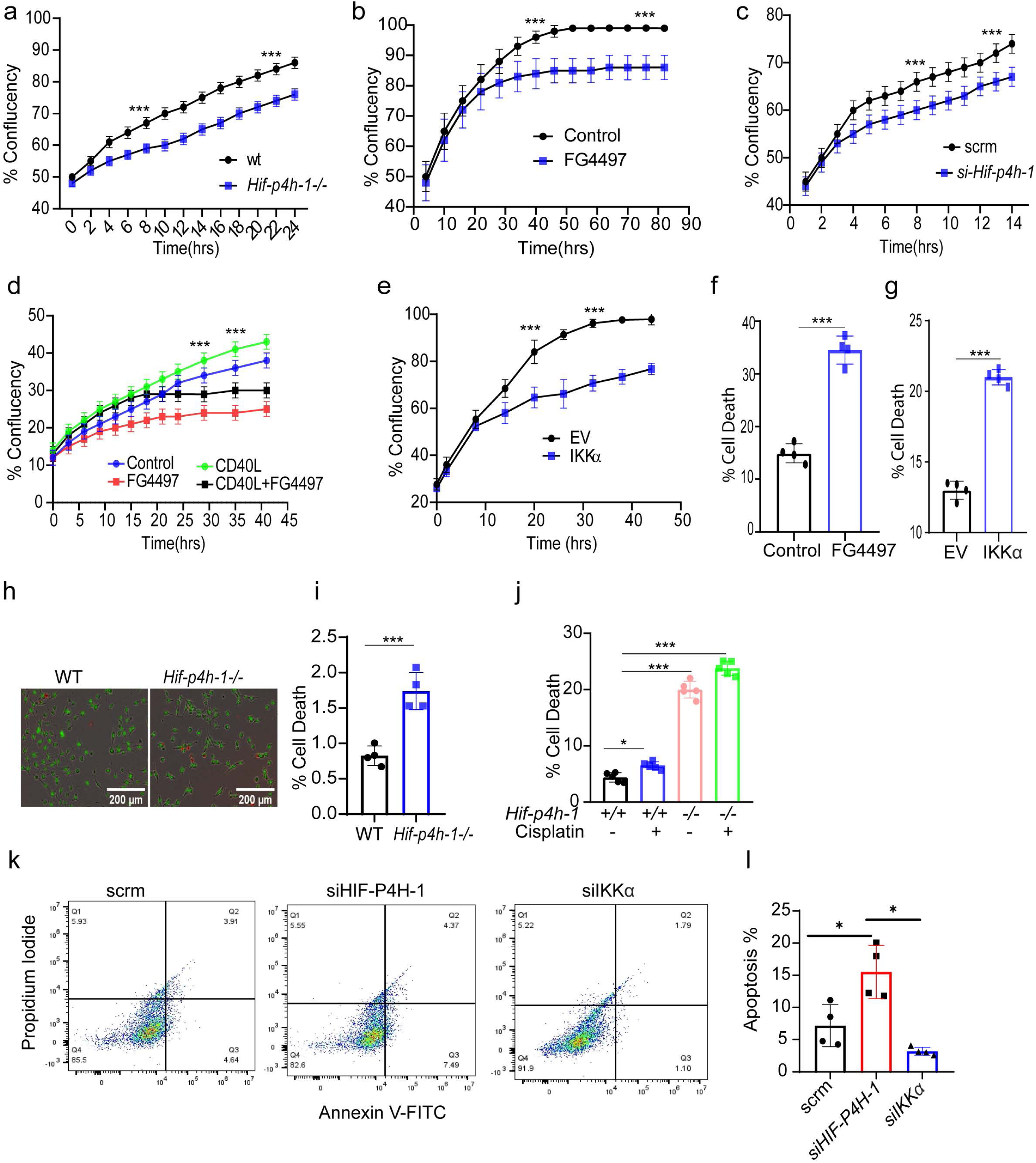
Inhibition of HIF-P4H-1 sensitizes cells to death by attenuating NF-κB2/p100 processing. (a) Proliferation of Hif-p4h-1⁻/⁻ and wild-type (WT) MEFs monitored using the IncuCyte™ Zoom live-cell imaging system (n = 8). (b) Proliferation of MDA-MB-231 cells treated with or without FG4497 (50 µM) assessed by IncuCyte™ Zoom (n = 8). (c) Proliferation of MDA-MB-231 cells transfected with HIF-P4H-1 siRNA, with confluence measured every 2 h using IncuCyte™ Zoom (n = 8). (d) MDA-MB-231 cells treated with FG4497 (50 µM) in the presence or absence of recombinant human CD40 ligand (hCD40L, 0.3 µg ml⁻¹), with proliferation determined by dynamic confluence measurements at 2-h intervals (n = 8). (e) Proliferation of MDA-MB-231 cells overexpressing full-length IKKα or empty vector measured by IncuCyte™ Zoom (n = 8). (f) Viability of M1 acute myeloid leukemia cells treated with FG4497 (50 µM, 24 h), assessed by 7-AAD staining and flow cytometry (n = 4). (g) Viability of MDA-MB-231 cells treated with FG4497 (50 µM) and staurosporine (2 µM, 24 h), measured by 7-AAD staining and flow cytometry (n = 4). (h, i) Live–dead staining of WT and Hif-p4h-1⁻/⁻ MEFs treated with cisplatin (100 µM) or left untreated, imaged using the IncuCyte™ Zoom system (n = 5). (k, l) Apoptosis of human medulloblastoma cells transfected with control siRNA or siRNAs targeting HIF-P4H-1 and/or IKKα was assessed by Annexin V staining and flow cytometry, with quantitative analysis shown (n = 4).

Analysis of cell death revealed significantly increased cell death following FG4497 treatment in M1 acute myeloid leukemia cells, as well as following HIF-P4H-1 silencing (Fig. 6f and Supplementary Fig. 7a) or IKKα overexpression (Fig. 6g) in MDA-MB-231 cells. This effect was further exacerbated by treatment with staurosporine, a widely used pan-kinase inhibitor that potently induces apoptosis, was included as a positive control to assess cellular sensitivity to apoptotic stress and further enhanced cell death under these conditions (Supplementary Fig. 7b). Similarly, Hif-p4h-1^-/-^ MEFs exhibited elevated levels of cell death under basal conditions, as assessed by TUNEL staining (Fig. 6h), and showed enhanced sensitivity to cisplatin-induced cell death (Fig. 6j). In agreement with these observations, gene set enrichment analysis demonstrated significant enrichment of cell death–associated transcriptional signatures in Hif-p4h-1^-/-^ MEFs (Supplementary Fig. 7c). Notably, genetic deletion of HIF-P4H-1 in a medulloblastoma cell model resulted in significantly increased apoptosis, as measured by Annexin V staining, whereas concomitant IKKα deletion abolished this effect (Fig. 6k–l). Similarly, in HCT116 cells treated with cisplatin, HIF-P4H-1 deletion markedly increased TUNEL-positive cells, while IKKα deletion significantly reduced cell death compared with both HIF-P4H-1–deficient and control-treated cells. These results indicate that excessive IKKα accumulation is a key contributor to the heightened apoptotic sensitivity observed following loss of HIF-P4H-1 (Fig. 7a, b).

**Figure 7:**
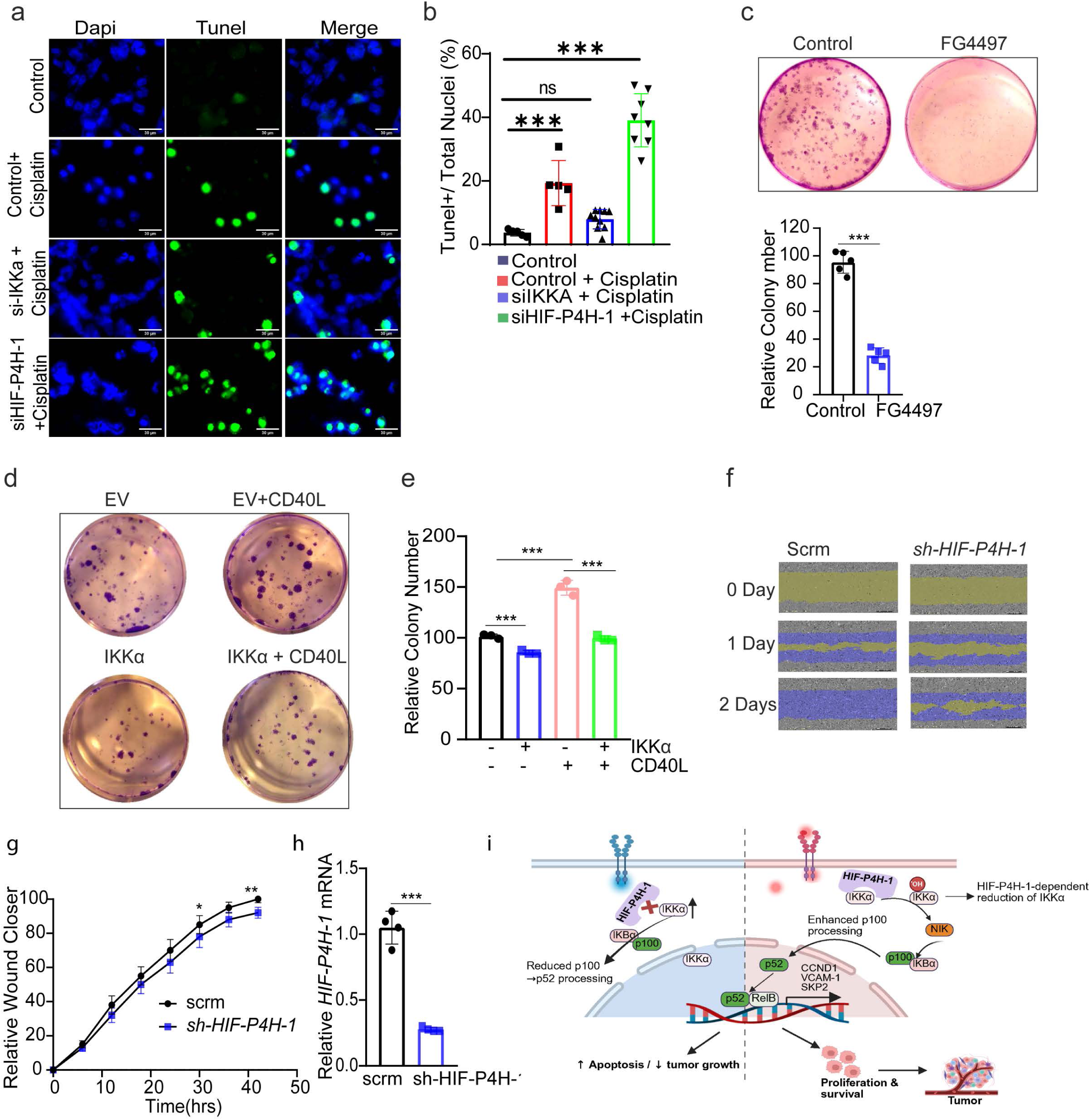
HIF-P4H-1–IKKα signaling promotes cancer cell survival, clonogenic growth, and migration. (a, b) Representative images (a) and quantification (b) of TUNEL-positive nuclei in HCT116 cells transfected with scrambled control siRNA, siHIF-P4H-1, or siIKKα, followed by cisplatin treatment (100 µM, 12 h). Nuclei were counterstained with DAPI. Scale bar, 30 µm. Sample sizes were n = 4 biological replicates for control siRNA and n = 8 biological replicates for siHIF-P4H-1 and siIKKα. Quantification is presented as the ratio of TUNEL-positive nuclei to total nuclei. (c) Representative images and quantification of colony formation in MDA-MB-231 cells treated with or without FG4497, with colony numbers quantified using ImageJ (n = 5 biologically independent replicates per group). (d, e) Colony formation of MDA-MB-231 cells overexpressing full-length IKKα following hCD40L stimulation over 2 weeks, visualized by crystal violet staining and quantified using ImageJ (n = 3 biologically independent replicates). (f–h) Cell migration analysis of shRNA-mediated HIF-P4H-1–depleted MDA-MB-231 cells monitored using the IncuCyte™ system, with HIF-P4H-1 knockdown efficiency confirmed by mRNA expression analysis (h). Data are presented as mean ± SD from n = 4–8 biologically independent experiments. (i) Schematic model illustrating HIF-P4H-1–dependent regulation of NF-κB2/p100 processing, cancer cell proliferation, and survival. For comparisons between two groups, Student’s t-test was used; for multiple comparisons, one-way ANOVA followed by Tukey’s post hoc test was applied. p < 0.05, *p < 0.01, **p < 0.001.

To determine whether the effects of HIF-P4H-1 inhibition extend beyond transformed cells, colony formation assays were performed in both MDA-MB-231 breast cancer cells and non-tumorigenic MDCK epithelial cells. Genetic ablation or pharmacological inhibition of HIF-P4H-1 markedly reduced clonogenic capacity in both cell types (Fig. 7c and Supplementary Fig. 8a). In parallel, CD40L-stimulated cells overexpressing IKKα exhibited significantly impaired colony formation (Fig. 7d, e), further supporting a functional antagonism between HIF-P4H-1 and IKKα signaling. Beyond effects on clonogenic growth, depletion of HIF-P4H-1 significantly suppressed migratory capacity in MDA-MB-231 cells, as demonstrated by migration assays (Fig. 7f–h). Collectively, these data indicate that HIF-P4H-1 promotes cell proliferation, survival, and migration by stabilizing IKKα and sustaining non-canonical NF-κB signaling. Accordingly, inhibition of HIF-P4H-1 attenuates NF-κB2/p100 processing, thereby sensitizing cells to apoptotic and genotoxic stress, as summarized in the proposed model (Fig. 7i). These findings establish HIF-P4H-1 as a key regulator of malignant phenotypes and highlight its potential as a therapeutic vulnerability in cancer.

### HIF-P4H-1 maintains NIK stability by modulating IKKα

NF-κB–inducing kinase (NIK), a central regulator of non-canonical NF-κB signaling, is normally maintained at low levels through constitutive degradation mediated by the TRAF–cIAP complex under basal conditions [24, 25]. Given that inhibition of HIF-P4H-1 reduced NF-κB2/p100 processing and that IKKα has been implicated in the regulation of NIK stability, we examined whether HIF-P4H-1 influences NIK protein abundance. Immunoblot analysis revealed a significant reduction in NIK levels in HIF-P4H-1-silenced cells compared to controls (Fig. 8a). Similarly, pharmacological inhibition with FG4497 lowered both endogenous and ectopically expressed NIK protein in MDA-MB-231 cells (Fig. 8b), an effect also observed after CD40L stimulation (Fig. 8c). Notably, reduced NIK protein was detected in TPA-treated in vivo skin samples and LPS-stimulated macrophages from *Hif-p4h-1^-/-^* mice, relative to respective controls (Fig. 8d, e). To assess whether this regulation involves IKKα, HEK293 cells were co-transfected with Flag-tagged IKKα and HA-tagged NIK. Overexpression of IKKα markedly decreased NIK protein abundance and was associated with reduced NF-κB2/p100 processing to p52, and lowered Cyclin D1 protein levels (Fig. 8f). Collectively, these data indicate that HIF-P4H-1 modulates NIK protein abundance indirectly through regulation of IKKα levels, thereby influencing non-canonical NF-κB signaling output, as summarized in the proposed graphical model in Fig. 8g.

**Figure 8:**
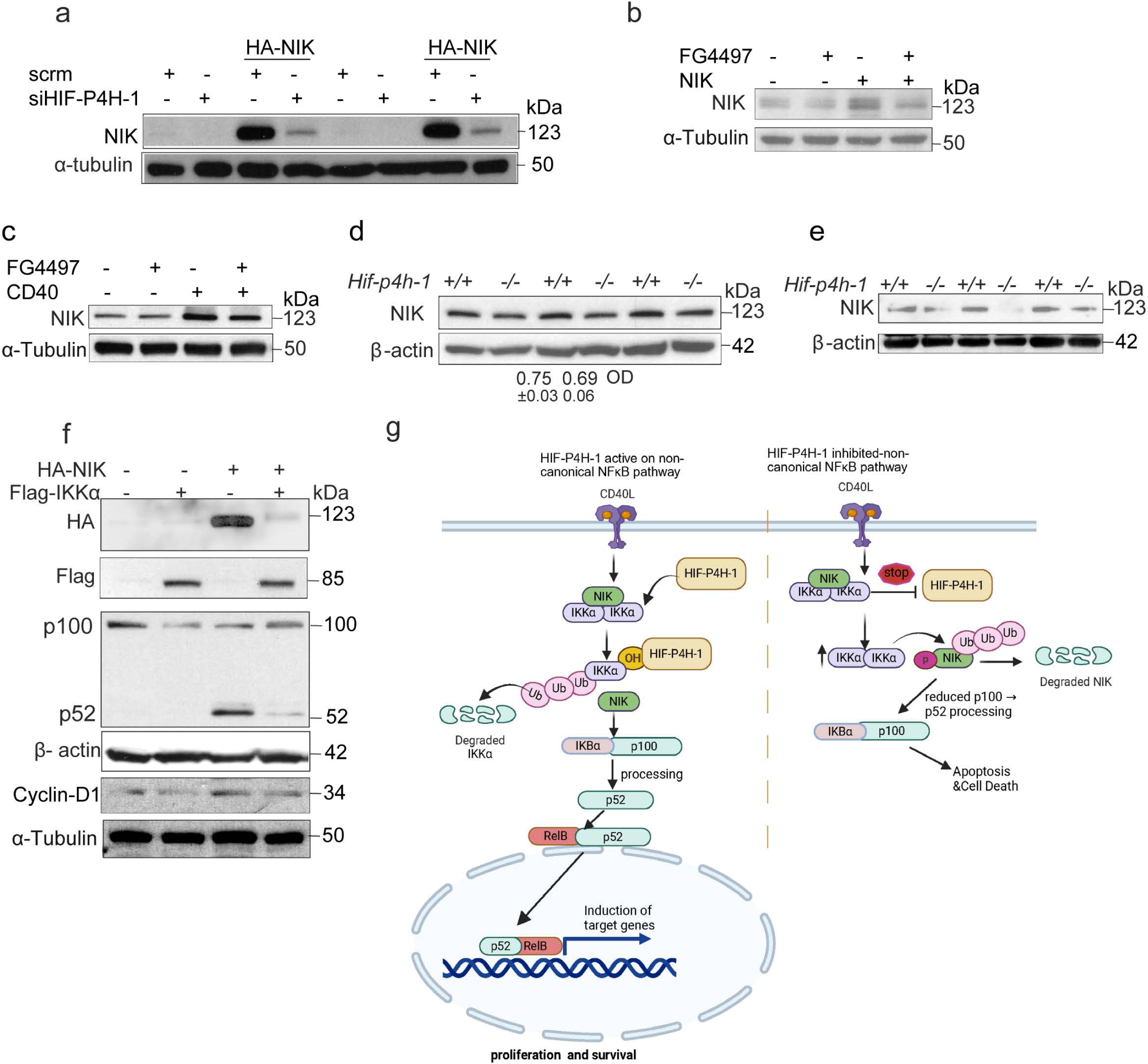
HIF-P4H-1 maintains NIK stability through modulation of IKKα. (a) siRNA-mediated knockdown of HIF-P4H-1 reduces NIK protein levels in HEK293 cells expressing HA-tagged NIK, as assessed by immunoblotting (n = 4 biologically independent replicates per group). (b) NIK protein levels in HA–NIK–transfected MDA-MB-231 cells treated with FG4497 (50 µM, 6 h), analyzed by immunoblotting (n = 3). (c) Immunoblot analysis of NIK protein levels in MDA-MB-231 cells stimulated with recombinant human CD40 ligand (hCD40L) in the presence or absence of FG4497 (n = 3). (d, e) Immunoblot analysis of endogenous NIK protein levels in Hif-p4h-1⁻/⁻ and wild-type macrophages (d) and in TPA-treated skin samples (e) (n = 4–5). (f) HEK293 cells co-expressing FLAG–IKKα and HA–NIK were analyzed by immunoblotting to assess the effect of IKKα on NIK abundance (n = 3). (g) Schematic model illustrating non-canonical NF-κB signaling, in which HIF-P4H-1 maintains NIK stability via hydroxylation-dependent regulation of IKKα, whereas loss or inhibition of HIF-P4H-1 promotes IKKα-mediated negative feedback and NIK destabilization.

## Discussion

In this study, we identify HIF-P4H-1 as a pivotal regulator of non-canonical NF-κB signaling in cancer cells, revealing a direct mechanistic link between oxygen sensing, post-translational modification of IKKα, and tumor cell survival. Specifically, our results demonstrate that inactivation or silencing of HIF-P4H-1 stabilizes IKKα, decreases processing of NF-κB2/p100 to p52, downregulates expression of survival genes, destabilizes NIK, and sensitizes cancer cells to apoptosis. These findings extend previous work on HIF-P4H-1 as a regulator of canonical NF-κB components and p53/FOXO3a signaling, as well as its broader roles in hypoxia adaptation and cyclin D1 control [14, 15, 17, 18, 22, 23, 26–28] and highlight the central role of the HIF-P4H-1–IKKα axis within the molecular stress-adaptation machinery of cancer. At the molecular level, we define a regulatory mechanism whereby HIF-P4H-1 interacts directly with IKKα and hydroxylates Pro367, thereby facilitating polyubiquitylation and proteasomal degradation of IKKα. Notably, this process operates independently of HIF-1α, distinguishing it from classical paradigms in which prolyl hydroxylases primarily modulate HIF-α stability [22, 27, 29]. Our identification of IKKα as a HIF-P4H-1 substrate expands the enzyme’s repertoire beyond previously characterized targets such as p53, FOXO3a, and Cep192 [14, 17–19], delineating a new oxygen-dependent control mechanism over the non-canonical NF-κB pathway.

The systems-level relevance of this pathway is underscored by transcriptomic and clinical data from TCGA cohorts. We observed a robust inverse correlation between EGLN2 (HIF-P4H-1) and CHUK (IKKα) expression in clear cell renal cell carcinoma (ccRCC), where high EGLN2 levels associate with advanced tumor stage and poor overall survival, while reduced CHUK expression correlates with unfavorable outcome. These observations support and extend existing evidence that hypoxia-regulated enzymes contribute to aggressive tumor phenotypes and adverse clinical behavior [20, 21]. In contrast, we did not observe significant survival effects of CHUK in breast cancer, indicating tissue-specific roles for non-canonical NF-κB signaling and providing impetus for further investigation into context-dependent mechanisms. To gain structural insight into how HIF-P4H-1 may regulate IKKα, we employed AlphaFold2-Multimer modeling to examine potential interactions of IKKα with HIF-P4H-1 and HIF-P4H-2. The models suggest that IKKα can transiently associate with the catalytic domains of HIF prolyl hydroxylases; however, Pro367 lies at a distance (>11 Å) from the Fe(II)-coordinating His–Asp–His triad, exceeding the optimal catalytic range (<5–7 Å). Interface confidence scores and spatial alignment metrics further indicated a recognition-competent but catalytically suboptimal interaction. These features likely reflect the absence of conformational flexibility and higher-order complexes in static models. Subsequent biochemical and cellular studies validated and refined these in silico predictions by demonstrating that HIF-P4H-1 directly interacts with and hydroxylates IKKα at Pro367 under physiological conditions, and that this modification is markedly reduced upon genetic or pharmacological inhibition of HIF-P4H-1. Mutating Pro367 to alanine in IKKα abolished hydroxylation and stabilized the protein, confirming the specificity of this post-translational modification and its role in targeting IKKα for ubiquitin-mediated proteolysis. Together, the complementary use of in silico and in vitro approaches provides a comprehensive mechanistic rationale for HIF-P4H-1–mediated regulation of IKKα and highlights both the utility and limitations of predictive protein–protein interaction modeling.

Our functional assays reveal a feedback mechanism whereby loss of HIF-P4H-1 activity promotes IKKα accumulation and nuclear localization, which in turn destabilizes NIK and suppresses inducible NF-κB2/p100 processing. The reduction of NIK observed in IKKα-accumulated cells aligns with its known negative-feedback loop, in which IKKα promotes NIK turnover [28], and provides a mechanistic explanation for the seemingly paradoxical finding that elevated IKKα is associated with reduced p52 generation. Attenuation of this pathway diminishes activation of key survival genes such as Cyclin D1, Bcl-xL, cIAP1, and VCAM-1, resulting in impaired proliferation, clonogenic growth, and increased apoptotic death across multiple cell models. These findings support a homeostatic role for HIF-P4H-1 in maintaining a dynamic balance between IKKα and NIK, thereby sustaining non-canonical NF-κB signaling competence under stress conditions. Therapeutically, our data indicate that both genetic deletion and pharmacological inhibition of HIF-P4H-1 (using FG4497) reproduce these cellular outcomes, validating HIF-P4H-1 as a promising molecular target. The recent clinical approval of pan-prolyl hydroxylase inhibitors for anemia [30], further supports the translational potential of this approach in oncology, where selective inhibition of HIF-P4H-1 could disrupt survival pathways in hypoxic tumor compartments and potentiate existing therapies. At the same time, these findings caution that systemic inhibition may compromise NF-κB–mediated survival responses that are crucial for normal tissue defense and homeostasis, underscoring the need for cell-type and context-specific targeting strategies.

Despite these advances, our study has limitations. Most mechanistic experiments were conducted in selected cell lines and in defined in vivo settings, which may not capture the full heterogeneity of human tumors or the complexity of the tumor microenvironment. The impact of HIF-P4H-1 inhibition on non-malignant tissues and immune function remains uncharacterized; given the broad physiological influence of NF-κB, this warrants careful investigation. Moreover, potential crosstalk between altered IKKα/NIK balance and other signaling networks, including canonical NF-κB and additional hypoxia-responsive pathways, was not addressed here and represents an important avenue for future work. Our findings are distinct from previous studies in which prolyl hydroxylases, particularly HIF-P4H-1, have been characterized as regulators of canonical NF-κB pathway components via hydroxylation-dependent control of protein stability [14, 17, 26]. To date, direct modulation of the non-canonical pathway by a HIF-P4H-1/IKKα axis has not been reported. Prior work by Razani et al.[28] described negative-feedback loops involving IKKα and NIK, but did not integrate oxygen-dependent regulation.

Studies focusing on HIF-P4H family members and substrates such as p53, FOXO3a, and Cep192 [14, 17–19] have hinted at a broad substrate scope for prolyl hydroxylases. Our current data directly connect oxygen-sensing HIF-P4H-1 activity to non-canonical NF-κB signaling within the broader context of NF-κB pathways implicated in tumor progression and treatment resistance [7, 10], and denote a tissue-and context-specific impact, as the inverse correlation between EGLN2 and CHUK in ccRCC is consistent with earlier work on cyclin D1 regulation in breast cancer [19] but reveals distinct survival associations. Notably, while our findings support a tumor-promoting role for HIF-P4H-1 in ccRCC, some clinical studies have identified HIF-1α expression as a favorable prognostic marker in renal cancer [20], pointing to isoform-specific and context-dependent biological effects. In addition, although pan-HIF-P4H inhibitors display antiproliferative effects in vitro [31], their systemic administration carries the risk of off-target or adverse effects, further emphasizing the need for precisely targeted and context-dependent therapeutic strategies.

### Future Perspectives

Future validation of this regulatory axis in primary patient-derived samples and orthotopic animal models will be essential to strengthen translational relevance. Comprehensive molecular profiling of HIF-P4H-1 inhibition within the tumor microenvironment, including its effects on immune, stromal, and metabolic networks, should be prioritized. Integrating multi-omics approaches with targeted HIF-P4H-1 inhibition and established therapeutic agents may reveal synergistic vulnerabilities and refine risk–benefit considerations. Additionally, delineating tissue-and context-specific determinants of HIF-P4H-1–IKKα–NF-κB2/p52 regulation may uncover new opportunities for precision oncology interventions.

### Limitation

A primary limitation of this study is the absence of a dedicated xenograft or orthotopic tumor model directly assessing how modulation of the HIF-P4H-1–IKKα axis influences tumor growth and therapy response in vivo. Nevertheless, prior xenograft studies have already established the pro-tumorigenic function of EGLN2/HIF-P4H-1 in vivo, including enhanced tumor growth and cyclin D1 induction, and our work now provides the mechanistic foundation for these observations by identifying IKKα as a hydroxylation-dependent substrate governing non-canonical NF-κB signaling [23]. Moreover, our analyses of patient tissues (ccRCC and glioblastoma), together with functional assays and in vivo murine skin models, provide convergent evidence supporting the biological relevance of this pathway. Collectively, these data strengthen the rationale for future tumor-focused in vivo studies specifically interrogating this regulatory mechanism.

## Conclusion

In summary, this study identifies a previously unrecognized, oxygen-dependent regulatory layer governing non-canonical NF-κB signaling through HIF-P4H-1–mediated hydroxylation of IKKα. By demonstrating that HIF-P4H-1 stabilizes NIK and thereby promotes p100 processing and p52-dependent transcription, we reveal a mechanistic link connecting hypoxia sensing to tumor-associated NF-κB activity. These findings lay the groundwork for therapeutically targeting HIF-P4H-1 as a strategy to modulate non-canonical NF-κB signaling in cancer and underscore the need for continued mechanistic and translational studies to ensure safe and effective clinical development.

## Methods

### Cell Lines and Rationale for Selection

A diverse panel of human and murine cell lines was used to investigate the role of HIF-P4H-1 in regulating IKKα stability and non-canonical NF-κB signaling across multiple tumor contexts. MDA-MB-231 triple-negative breast cancer cells were selected as a well-established model exhibiting robust non-canonical NF-κB pathway activity and were used for mechanistic, signaling, and functional studies. HCT116 HIF1A-/- cells and their matched wild-type counterparts were included to evaluate whether regulation of IKKα by HIF-P4H-1 occurs independently of HIF1α and to assess apoptosis in response to DNA-damage stress. HEK293 cells were used for co-immunoprecipitation, hydroxylation, and ubiquitination assays owing to their high transfection efficiency and suitability for biochemical analyses. To examine whether the HIF-P4H-1–IKKα axis extends beyond solid tumors, M1 murine acute myeloid leukemia cells were included for viability analyses under pharmacological inhibition of HIF-P4H-1. In addition, human medulloblastoma cell lines representing low-and high-aggressive phenotypes were incorporated to evaluate tumor aggressiveness–associated differences in HIF-P4H-1 and IKKα protein abundance and to assess apoptosis following genetic perturbation of these factors. Murine Hif-p4h-1^-/-^and Hif-p4h-3^-/-^mouse embryonic fibroblasts, as well as primary macrophages and TPA-treated mouse skin samples, were used to validate findings under physiological conditions and to examine the effects of genetic loss of HIF-P4H isoforms on IKKα stability and NF-κB2/p100 processing. All cell lines were routinely tested and confirmed to be free of mycoplasma contamination.

### Cell Culture and Transfections

Human embryonic kidney (HEK293) and human colon carcinoma HCT116 (*HIF1α ^+/+^* and HCT116 *HIF1α ^-/-^*) cells were maintained in DMEM medium (Invitrogen). MDA-MB-231 human breast adenocarcinoma cells were maintained in RPMI-1640 medium (Thermo Scientific). The myeloid leukemia cell lines M1 and TIB-49 were cultured in suspension in RPMI-1640. *Hif-p4h-1^-/-^*and WT gender-matched mouse embryonic fibroblasts (MEFs) were isolated at 14.5 embryonic days and maintained in DMEM medium [17]. Murine peritoneal macrophages were isolated from 1-year-old female *Hif-p4h-1^-/-^*and wt mice as described previously [32]. All cell culture media were supplemented with 10% FBS (Gibco) and 1% penicillin/streptomycin (Gibco) at 37°C in a humidified chamber with 5% CO₂. All cell lines tested negative for mycoplasma contamination and were not listed as misidentified by ICLAC or NCBI Biosample databases. Cells were treated as indicated with FG4497 (Fibrogen, 50 μM, 6 h), MG132 (Calbiochem, 10 to 30 μM), cisplatin (Santa Cruz, 100 μM), LPS (Sigma, 200 ng/mL), or recombinant human CD40L (R&D Systems, 0.3 μg/mL, overnight). A mammalian pcDNA plasmid encoding recombinant human HIF-P4H-1 with a C-terminal V5-His tag was generated as previously described for HIF-P4Hs [33]. pCR-Flag-IKKα and pCMV4-NIK-HA vectors were obtained from Addgene (#15467 and #27554). Plasmid transfections were performed using FuGENE® HD (Promega) or Lipofectamine LTX (Thermo Fisher) according to the manufacturer’s protocols, in cells at 70–80% confluence. Small interfering RNAs targeting HIF-P4H-1 (Santa Cruz, sc-45616 and Silencer Select siRNAs EGLN2 #AM16708 ) or control siRNA (sc-37007) were transfected using RNAiMAX (Invitrogen).

### Western Blotting

Whole-cell extracts were prepared using lysis buffer (50 mM Tris-HCl, pH 8.0, 150 mM NaCl, and 1% Triton X-100) supplemented with phosphatase (PhosSTOP, Roche) and protease (Complete, Roche) inhibitors. Cytosolic and nuclear fractions were obtained with the PARIS™ kit (Life Technologies) according to the manufacturer’s instructions. Proteins were resolved by SDS–PAGE, transferred to PVDF membranes, and probed with the indicated primary antibodies. The following antibodies and dilutions were used: anti-IKKα (Novus Biologicals, #14A231; 1:500), cyclin D1 (Fisher Scientific, #RB010P0; 1:5000), anti-FLAG M2 (Sigma, #F1804; 1:1000), anti-NFκB2 (Cell Signaling, #3017 or #4882; 1:500), anti-V5 (Life Technologies, #R96125; 1:500), anti-hydroxyproline (Abcam, #ab37067; 1:300), anti-HA (Sigma; 1:1000), anti-NIK (Cell Signaling, #4994; 1:500), anti-Bcl-XL (Novus Biologicals, #NB100-56104; 1:1000), anti-β-actin (Novus Biologicals, #NB600-501; 1:20,000), and anti-α-tubulin (Sigma, #T6199; 1:20,000). Secondary antibodies included HRP-conjugated goat anti-rabbit IgG (Dako; 1:10,000) and HRP-conjugated goat anti-mouse IgG (Dako; 1:10,000). Signals were visualized using enhanced chemiluminescence (Pierce™ ECL Western Blotting Substrate).

### Structural Modeling with AlphaFold2-Multimer

Protein–peptide complex predictions were generated using ColabFold (AlphaFold2-Multimer v3) [34]. A 21-residue IKKα peptide (residues 356–377, centered on Pro367) was modeled against the catalytic domains of HIF-P4H-2 (residues 181–426) and HIF-P4H-1 (residues 278–407). For mutant models, Pro367 was substituted with Ala. Structural metrics, including distances between Pro367/Ala367 and the catalytic His–Asp–His triad, root mean square deviation (RMSD), and solvent-accessible surface area (SASA), were quantified in PyMOL. Model quality parameters (pTM, iPTM, actifpTM, pLDDT, and pAE) were obtained from ColabFold outputs and analyzed.

### Co-immunoprecipitation

Cells were lysed in lysis buffer, and lysates were sonicated. A total of 500 μg of protein was pre-cleared with 30 μl Protein A/G Plus agarose beads (Santa Cruz Biotechnology). For FLAG immunoprecipitation, 50 μl Anti-FLAG M2 affinity gel (Sigma-Aldrich) was added to pre-cleared lysates and incubated for 2 h at 4 °C. Beads were washed three times with wash buffer (0.5 M Tris-HCl, 1.5 M NaCl, pH 7.4). Bound proteins were eluted with 300 ng/μl FLAG peptide (Sigma-Aldrich) by incubation for 45 min at 4 °C. For V5 immunoprecipitation, 3 μl anti-V5 antibody (Life Technologies) was incubated with pre-cleared lysates for 2 h at 4 °C, followed by addition of 30 μl Protein A/G Plus agarose beads (Santa Cruz Biotechnology) and incubation for 1 h at 4 °C. Beads were washed three times with wash buffer, and proteins were eluted using a soft elution protocol [35]. Eluted proteins were analyzed by Western blotting using the indicated antibodies.

### Immunoprecipitation for LC–MS, Radiochemical, and Western Blot Analyses of 4-Hydroxyproline

MDA-MB-231 cells were transfected with FLAG-tagged IKKα and treated with MG132 for 6 h. Cell extracts were prepared using lysis buffer as described above, and 5 mg of total protein was pre-cleared before immunoprecipitation. FLAG-tagged proteins were captured with 50 μl Anti-FLAG M2 affinity gel (Sigma-Aldrich) and incubated overnight at 4 °C with rotation. Beads were washed three times with wash buffer (0.5 M Tris-HCl, 1.5 M NaCl, pH 7.4). Bound proteins were eluted with 300 ng/μl FLAG peptide (Sigma-Aldrich) in TBS and incubated for 45 min at 4 °C. Eluates were resolved by SDS–PAGE and subjected to LC–MS analysis. For radiochemical detection of 4-hydroxyproline, MDA-MB-231 cells were transfected with FLAG-tagged IKKα and incubated for 48 h with [2,3,4,5-3H]proline (25 μCi/ml). Cells were treated with 20 μM MG132 in the presence or absence of 50 μM FG4497 for 6h before lysis. FLAG-tagged IKKα was immunoprecipitated as described above, and the incorporation of 4-hydroxy[3H]proline was quantified using a radiochemical assay [36]. In parallel, FLAG-IKKα immunoprecipitates were analyzed by Western blotting with the indicated antibodies.

### Liquid Chromatography–Mass Spectrometry (LC–MS) In-gel digestion

FLAG–IKKα-containing regions of the SDS–PAGE gel were excised and cut into ∼1 mm slices. Gel pieces were destained with three 5-mins washes in 50 mM ammonium bicarbonate/40% acetonitrile, followed by a 5-min wash in trypsin buffer (40 mM ammonium bicarbonate, 9% acetonitrile). Proteins were digested by adding 5 μl proteomics-grade trypsin (20 ng/μl; Sigma) in trypsin buffer and incubating for 20 min, followed by addition of 15 μl trypsin buffer. Digestion proceeded overnight at 37 °C. Peptides were sequentially extracted with 0.1% trifluoroacetic acid (TFA) and TA30 (30% acetonitrile, 0.1% TFA). Combined extracts were dried in a SpeedVac and reconstituted in 20 μl 0.2% TFA prior to LC–MS analysis.

### LC–MS analysis (Orbitrap Fusion Lumos)

Peptides were analyzed on an Easy-nLC 1000 (Thermo) coupled to a Fusion Lumos Tribrid mass spectrometer (Thermo). Samples were first trapped on a Symmetry C18 trap column (0.18 × 20 mm; Waters) and separated on a BEH C18 analytical column (75 μm × 150 mm, 1.7 μm, 130 Å; Waters) using a linear gradient from 97% solvent A (0.1% formic acid in water) to 35% solvent B (0.1% formic acid in acetonitrile) over 90 min at 0.3 μl/min. Data were acquired in 3-sec cycles: full MS scans in the Orbitrap (resolution 120,000; AGC 4e5; max injection time 50 ms), followed by MS/MS of multiply charged precursors (>5e4 intensity) using HCD (30% CE) and CID (35% CE, 10 ms activation, q = 0.25). Isolation width was 1.6 Da with a 21-sec dynamic exclusion. Fragment ions (up to 5e4) were accumulated for 200 ms and analyzed in the Orbitrap (resolution 30,000). A second acquisition method collected MS/MS in the ion trap (rapid mode; threshold 1e4) for enhanced sensitivity.

### LC–MS analysis (Synapt G2 Q-TOF)

Parallel analyses were performed on a nanoAcquity UPLC system (Waters) coupled to a Synapt G2 Q-TOF mass spectrometer (Waters). Samples were trapped on a Symmetry C18 trap column (0.18 × 20 mm) for 5 min (flow 5 μl/min) and separated on a PicoFrit column (75 μm × 150 mm; New Objective) packed with BEH C18 resin (1.7 μm, 130 Å). Peptides were eluted with a linear gradient from 3% to 40% solvent B over 80 min (solvent A: 0.1% formic acid in water; solvent B: 0.1% formic acid in acetonitrile) at 0.3 μl/min. The instrument operated in MSE mode (data-independent acquisition), alternating between low-energy (5 V) and elevated-energy (18–40 V ramp) scans (0.5 s per scan).

### Data processing

Orbitrap data were analyzed in Proteome Discoverer 2.2 (Thermo) using Sequest. Spectra were searched against the SwissProt human database (release 2017-07-05; 9605 entries), supplemented with a custom IKKα sequence (O15111, P367A mutation). Search parameters included: precursor mass tolerance 10 ppm, fragment tolerance 0.02 Da (Orbitrap) or 0.6 Da (ion trap), trypsin specificity with up to two missed cleavages, fixed carbamidomethylation (C), and variable modifications (oxidation: M, P, A, K; deamidation: N, Q; N-terminal acetylation). False discovery rate (FDR) was controlled at 1% with Percolator. Label-free quantification used the Minora feature detector (2-min RT alignment, precursor intensity normalized to total peptide abundance). Synapt data were processed in PLGS 2.5 (Waters) with automatic determination of mass tolerance and peak width. Spectra were searched against SwissProt (human) allowing phosphorylation (S, T, Y), oxidation (M, W, P), N-terminal acetylation, and deamidation (N, Q). FDR was set to 4% liquid Chromatography–Mass Spectrometry (LC–MS)

### TCGA Data Extraction and Analysis

RNA-sequencing gene expression and matched clinical data for EGLN2 (HIF-P4H-1) and CHUK (IKKα) were obtained from The Cancer Genome Atlas (TCGA) KIRC cohort using the UCSC Xena browser (https://xenabrowser.net/) and GEPIA2 (http://gepia2.cancer-pku.cn/). Only samples with available tumor stage and survival outcome data were included. For differential gene expression analyses, default cutoffs in GEPIA2 were applied: log₂ fold-change (|log₂FC| > 1) and adjusted P-value (< 0.01). Expression comparisons, Pearson correlation analysis, Kaplan–Meier survival analysis, and tumor stage association were performed using built-in tools on the aforementioned platforms. All statistical outputs, including correlation coefficients, p-values, hazard ratios, and survival curves, were generated through these web-based interfaces. Coexpression analysis of HIF-P4H-1 and CHUK (IKKα) was conducted using cBioPortal for Cancer Genomics (https://www.cbioportal.org/), with Pearson correlation coefficients computed via the portal’s analysis module. Graphical visualization of correlation results was performed using GraphPad Prism (version 8.3.0). Publicly available TCGA datasets were utilized for all analyses.

### Quantitative Real-Time PCR (qRT-PCR)

Total RNA was extracted from cells using the E.Z.N.A. Total RNA Kit I (Omega Bio-Tek) according to the manufacturer’s instructions. One microgram of RNA was reverse transcribed into cDNA using the iScript Reverse Transcription Supermix (Bio-Rad). The resulting cDNA was diluted 1:10 before analysis. qRT-PCR was performed with iTaq Universal SYBR Green Supermix (Bio-Rad) on a CFX96 Real-Time PCR Detection System (Bio-Rad). Gene expression levels were normalized to β-actin mRNA. Primer sequences are listed in Supplementary Table 1.

### Luciferase Reporter Assays

Luciferase activity was measured using the Dual-Luciferase Reporter Assay System (Promega) according to the manufacturer’s instructions. Cells were co-transfected with κB-TATA–luciferase and a constitutively active Renilla luciferase control (pRL-TK). After 24–48 h, cells were lysed in Passive Lysis Buffer (Promega), and luciferase activity was measured. Renilla luciferase was used to normalize κB-TATA reporter activity. All experiments were performed in triplicate, and statistical significance was defined as P < 0.05.

### Cell Viability Assays

*Hif-p4h-1^-/-^*and wild-type MEFs were seeded in 12-well plates and stained using the Live/Dead Staining Kit (BioVision). Images were acquired with the IncuCyte ZOOM™ live-cell imaging system (Essen BioScience, MI, USA). For flow cytometry analysis, cells were plated in 3 cm² plates at a density of 5 × 10^5 cells per plate and treated with 50 μM FG4497, 100 μM cisplatin (Santa Cruz, #sc-200896), 2 μM staurosporine (Santa Cruz, #sc-3510), or transfected with full-length IKKα or empty vector. Following treatment, cells were washed with PBS, stained with saturating concentrations of 7-AAD, and analyzed using a FACSCalibur flow cytometer (BD Biosciences). Medublastoma cell lines D283 were transfected with HIF-P4H-1 siRNA or IKKα siRNA. After 72 hours, cells were stained with annexin V and propidium iodide (BD Bioscience) and analyzed by flow cytometry. The raw data were processed by FlowJo (version 10).

### Colony Formation Assays

For colony formation assays, 500–2,000 cells were seeded in 12-well plates and treated with 10 ng/mL TGFβ, 50 μM FG4497, or transfected with IKKα. After 10 days, colonies were stained with crystal violet (Sigma) and quantified using ImageJ software.

### Cell Proliferation Assays

Cell proliferation was monitored using the IncuCyte ZOOM™ live-cell imaging system (Essen BioScience, MI, USA). Cells were seeded in 12-well plates at a density of 5 × 10^4 cells per well and treated with 0.3 μg/mL hCD40L, with or without 50 μM FG4497, for 24–48 h. Images were acquired every 2 h, and proliferation was quantified using the accompanying software. HIF-P4H-1 siRNA or IKKα siRNA-transfected medulloblastoma cell line was stained intracellularly with anti-HIF-P4H-1 (Abcam) or IKKα antibodies (Thermofisher) and labeled with secondary anti-rabbit or anti-mouse antibodies (Thermofisher).

### Medulloblastoma cell model

#### Tumor collection and dissociation

Tumor specimens were collected following informed patient consent and in accordance with Dana–Farber/Harvard Cancer Center Institutional Review Board protocol #10-417. Full details of the protocol and associated consent documentation are available through the ODQ Neuro-Oncology database. Tissue dissection and sample acquisition during neurosurgical procedures were performed at the discretion of the operating physicians. All specimens were initially processed by the pathology department, which determined the amount of tissue required for clinical diagnostics. Excess tissue not needed for patient care was released for research use.

#### Immunofluorescence and histological analysis

Paraffin-embedded tumor tissues were sectioned at 5 µm and stained with hematoxylin and eosin (H&E) for histopathological evaluation. Images were acquired using an inverted light microscope with 20× and 40× objectives (Zeiss Axio Observer Z1; Carl Zeiss Microscopy, Thornwood, NY). For immunofluorescence analysis, fresh tumor tissues were dissected and fixed in 4% paraformaldehyde (PFA) for 24 h, followed by cryoprotection in 30% sucrose for 72 h. Samples were embedded in optimum cutting temperature compound (Tissue-Plus O.C.T.; Fisher Healthcare, Thermo Fisher Scientific) and cryosectioned at 5 µm thickness. Paraffin sections were deparaffinized in xylene and rehydrated through a graded ethanol series to water. Before antigen retrieval, slides were fixed in 10% neutral-buffered formalin for 30 min. Heat-induced antigen retrieval was performed using retrieval buffer (10 mM Tris-HCl, 1 mM EDTA, 10% glycerol; pH 9) at 110 °C for 17 min (∼4–5 psi). Sections were allowed to cool to room temperature and washed once in PBS for 15 min. Slides were blocked with 2.5% goat serum (Vector Laboratories, S-1012) for 90 min and incubated overnight at 4 °C with primary antibodies against HIF-P4H-1 (Invitrogen, OTI7H3) and IKKα (Invitrogen, #23GB765). After washing, sections were incubated for 60 min at room temperature with fluorophore-conjugated secondary antibodies (1:1000; Alexa Fluor 488 donkey anti-goat, cat. #A11055; Alexa Fluor 594 goat anti-rabbit, cat. #A11012 and A11037; Invitrogen, Thermo Fisher Scientific) with gentle agitation.Nuclei were counterstained with DAPI, and coverslips were mounted using ProLong Gold Antifade Mountant (P36931; Life Technologies). Images were acquired and processed using a confocal laser-scanning microscope with a 20× objective (Zeiss LSM 880 with Airyscan; Carl Zeiss Microscopy).

#### Microarray Analysis

Microarray data (GEO accession: GSE83535) were previously described [17]. Affymetrix CEL files were analyzed using dChip 2010_01 software. CEL files were normalized, and differential gene expression between wild-type (baseline [B]) and *Hif-p4h-1^−/−^* (experimental [E]) MEFs under normoxia and hypoxia was determined using the comparison function of dChip. Filtering criteria were: (a) fold change ≥1.5 (E/B or B/E), (b) lower confidence bound of fold change = 100, and (c) p < 0.05 by paired t-test. Differentially expressed genes were annotated using dChip’s gene function enrichment module, and functional classifications were further verified with DAVID Bioinformatics Resources. Gene set enrichment analysis (GSEA) was performed using GSEA tools (http://software.broadinstitute.org/gsea/index.jsp

#### Statistical Analysis

All statistical analyses were conducted using GraphPad Prism software (version 8.3.0, GraphPad, San Diego, CA). Data are expressed as mean ± standard error of the mean (SEM). The normality of each dataset was evaluated using the Shapiro-Wilk test. For comparison between two groups with normally distributed data, an unpaired two-tailed Student t-test was applied. For comparisons among multiple groups, a one-way ANOVA followed by Tukey’s multiple comparisons test or a two-way ANOVA with Bonferroni post hoc test was used, depending on the number of experimental variables. A p-value of < 0.05 was considered statistically significant (*, P < 0.05; **, P < 0.01; ***, P < 0.001).

## DATA Availability

The microarray datasets analyzed in this study were previously published and are publicly available in the Gene Expression Omnibus (GEO) under accession number GSE83535 (https://www.ncbi.nlm.nih.gov/geo/). All original, uncropped Western blot images supporting the findings of this study are provided in the Supplementary Information.

## Acknowledgments

We are grateful to M. Siurua, R. Juntunen, A. Kokko, E. Lehtimäki, and R. Polojärvi for excellent technical assistance, and to Prof. P. Koivunen for providing the *HIF1A^-/-^*and wild-type HCT116 cells. We also acknowledge the Biocenter Oulu Transgenic Core Facility, supported by the University of Oulu and Biocenter Finland. This work was funded by the Academy of Finland Center of Excellence (2012–2017, Grant 251314), the S. Jusélius Foundation, the Jane and Aatos Erkko Foundation, and FibroGen, Inc. Schematic illustrations in the figures were created using BioRender.com.

## Author Contributions

JM and KU conceived the study and designed the research strategy. JM and JMM supervised the project. JMM generated the *Hif-p4h-1* and *Hif-p4h-3* mouse lines. KU performed all experiments presented in Figures 1–8 and wrote the manuscript. MR and AG carried out the medulloblastoma tissue analyses. FQ performed the AlphaFold2 structural modeling. J.Z. and Q.Z. characterized and provided the ccRCC tumor samples. VI assisted with flow cytometry analyses, and UB performed the LC–MS/MS experiments. RW, JMM, and JM provided scientific input and overall support throughout the study.

## Ethics declarations

### Competing Interests

J.M. has a collaboration agreement with FibroGen Inc., approved by the Academy of Finland and the University of Oulu. J.M. owns equity in the company, which supports research in the J.M. laboratory. Other authors declare no competing interests.

### Ethics Approval

All human tumor sample–related studies were approved by the Institutional Review Boards (IRB) of the University of Texas Southwestern Medical Center and the University of North Carolina at Chapel Hill. All specimens were fully de-identified before analysis and processed in accordance with approved protocols and ethical guidelines. Informed consent was obtained from all subjects or waived by the IRBs as appropriate.

Animal experiments were approved by the Animal Experiment Board of Finland and conducted under the EU Directive 86/609/EEC, the European Convention ETS123, and Finnish national legislation. All procedures followed the recommendations of the Federation of European Laboratory Animal Science Associations (FELASA) and Finnish and EU legislation.

## Notes

### Summary of Updates

All authors are included in this submission.

## References

1. Khan, S.U., et al., Unveiling the mechanisms and challenges of cancer drug resistance. Cell Commun Signal, 2024. 22(1): p. 109.

2. Vijayakumar, S., et al., Drug resistance in human cancers -Mechanisms and implications. Life Sci, 2024. 352: p. 122907.

3. Cree, I.A. and P. Charlton, Molecular chess? Hallmarks of anti-cancer drug resistance. BMC Cancer, 2017. 17(1): p. 10.

4. Li, Y., et al., Inhibition of NF-kappaB signaling unveils novel strategies to overcome drug resistance in cancers. Drug Resist Updat, 2024. 73: p. 101042.

5. Mao, H., X. Zhao, and S.C. Sun, NF-kappaB in inflammation and cancer. Cell Mol Immunol, 2025. 22(8): p. 811–839.

6. Hayden, M.S. and S. Ghosh, Shared principles in NF-kappaB signaling. Cell, 2008. 132(3): p. 344–62.

7. Sun, S.C., Non-canonical NF-kappaB signaling pathway. Cell Res, 2011. 21(1): p. 71–85.

8. Sun, S.C. and S.C. Ley, New insights into NF-kappaB regulation and function. Trends Immunol, 2008. 29(10): p. 469–78.

9. Perkins, N.D., NF-kappaB: tumor promoter or suppressor? Trends Cell Biol, 2004. 14(2): p. 64–9.

10. Demchenko, Y.N., et al., Classical and/or alternative NF-kappaB pathway activation in multiple myeloma. Blood, 2010. 115(17): p. 3541–52.

11. Keats, J.J., et al., Promiscuous mutations activate the noncanonical NF-kappaB pathway in multiple myeloma. Cancer Cell, 2007. 12(2): p. 131–44.

12. Solt, L.A. and M.J. May, The IkappaB kinase complex: master regulator of NF-kappaB signaling. Immunol Res, 2008. 42(1-3): p. 3–18.

13. Sun, S.C., The noncanonical NF-kappaB pathway. Immunol Rev, 2012. 246(1): p. 125–40.

14. Cummins, E.P., et al., Prolyl hydroxylase-1 negatively regulates IkappaB kinase-beta, giving insight into hypoxia-induced NFkappaB activity. Proc Natl Acad Sci U S A, 2006. 103(48): p. 18154–9.

15. Rius, J., et al., NF-kappaB links innate immunity to the hypoxic response through transcriptional regulation of HIF-1alpha. Nature, 2008. 453(7196): p. 807–11.

16. Walmsley, S.R., et al., Hypoxia-induced neutrophil survival is mediated by HIF-1alpha-dependent NF-kappaB activity. J Exp Med, 2005. 201(1): p. 105–15.

17. Ullah, K., et al., Hypoxia-inducible factor prolyl-4-hydroxylase-1 is a convergent point in the reciprocal negative regulation of NF-kappaB and p53 signaling pathways. Sci Rep, 2017. 7(1): p. 17220.

18. Moser, S.C., et al., PHD1 links cell-cycle progression to oxygen sensing through hydroxylation of the centrosomal protein Cep192. Dev Cell, 2013. 26(4): p. 381–92.

19. Zheng, X., et al., Prolyl hydroxylation by EglN2 destabilizes FOXO3a by blocking its interaction with the USP9x deubiquitinase. Genes Dev, 2014. 28(13): p. 1429–44.

20. Lidgren, A., et al., The expression of hypoxia-inducible factor 1alpha is a favorable independent prognostic factor in renal cell carcinoma. Clin Cancer Res, 2005. 11(3): p. 1129–35.

21. Jiang, Y., et al., Temporal regulation of HIF-1 and NF-kappaB in hypoxic hepatocarcinoma cells. Oncotarget, 2015. 6(11): p. 9409–19.

22. Myllyharju, J., Prolyl 4-hydroxylases, master regulators of the hypoxia response. Acta Physiol (Oxf), 2013. 208(2): p. 148–65.

23. Zhang, Q., et al., Control of cyclin D1 and breast tumorigenesis by the EglN2 prolyl hydroxylase. Cancer Cell, 2009. 16(5): p. 413–24.

24. Sun, S.C., Controlling the fate of NIK: a central stage in noncanonical NF-kappaB signaling. Sci Signal, 2010. 3(123): p. pe18.

25. Rapino, F., et al., NIK is required for NF-kappaB-mediated induction of BAG3 upon inhibition of constitutive protein degradation pathways. Cell Death Dis, 2015. 6(3): p. e1692.

26. Cummins, E.P., et al., The hydroxylase inhibitor dimethyloxalylglycine is protective in a murine model of colitis. Gastroenterology, 2008. 134(1): p. 156–65.

27. Myllyharju, J. and P. Koivunen, Hypoxia-inducible factor prolyl 4-hydroxylases: common and specific roles. Biol Chem, 2013. 394(4): p. 435–48.

28. Razani, B., et al., Negative feedback in noncanonical NF-kappaB signaling modulates NIK stability through IKKalpha-mediated phosphorylation. Sci Signal, 2010. 3(123): p. ra41.

29. Fong, G.H. and K. Takeda, Role and regulation of prolyl hydroxylase domain proteins. Cell Death Differ, 2008. 15(4): p. 635–41.

30. Bartnicki, P., Hypoxia-Inducible Factor Prolyl Hydroxylase Inhibitors as a New Treatment Option for Anemia in Chronic Kidney Disease. Biomedicines, 2024. 12(8).

31. Tapio, J., et al., Contribution of HIF-P4H isoenzyme inhibition to metabolism indicates major beneficial effects being conveyed by HIF-P4H-2 antagonism. J Biol Chem, 2022. 298(8): p. 102222.

32. Ray, A. and B.N. Dittel, Isolation of mouse peritoneal cavity cells. J Vis Exp, 2010(35).

33. Hirsila, M., et al., Effect of desferrioxamine and metals on the hydroxylases in the oxygen sensing pathway. FASEB J, 2005. 19(10): p. 1308–10.

34. Mirdita, M., et al., ColabFold: making protein folding accessible to all. Nat Methods, 2022. 19(6): p. 679–682.

35. Antrobus, R. and G.H. Borner, Improved elution conditions for native co-immunoprecipitation. PLoS One, 2011. 6(3): p. e18218.

36. Koivunen, P., et al., The length of peptide substrates has a marked effect on hydroxylation by the hypoxia-inducible factor prolyl 4-hydroxylases. J Biol Chem, 2006. 281(39): p. 28712–20.

