## Supplementary Figures for "Inactivation of HIF-P4H-1 Stabilizes IKKα, Modulates Non-Canonical NF-κB Signaling, and Sensitizes Cancer Cells to Cell Death": Supplementary Figures 1-8.pdf

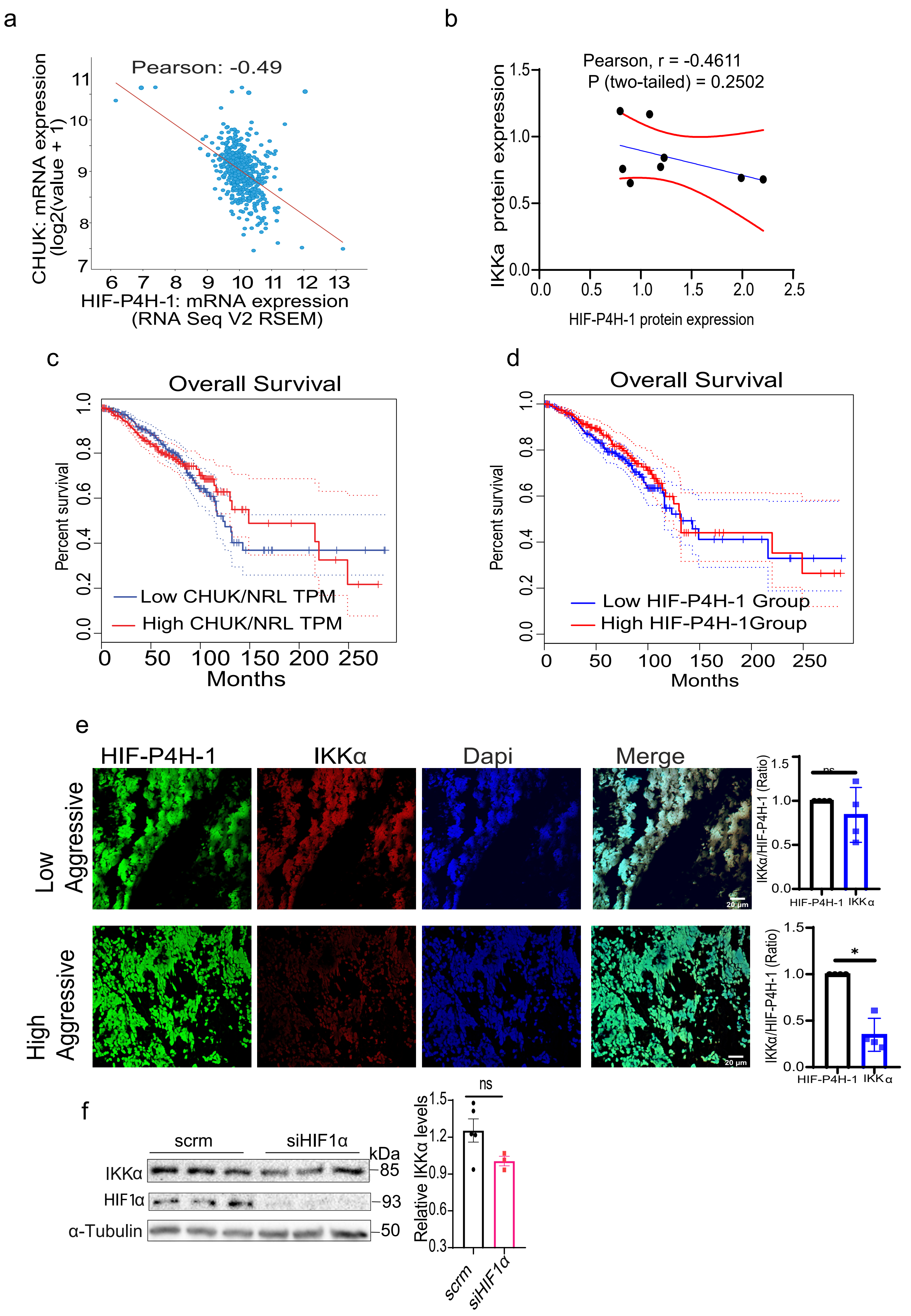

a
Pearson: -0.49
CHUK: mRNA expression (log2(value + 1))
HIF-P4H-1: mRNA expression (RNA Seq V2 RSEM)
b
Pearson, r = -0.4611
P (two-tailed) = 0.2502
IKKa protein expression
HIF-P4H-1 protein expression
c
Overall Survival
Percent survival
Low CHUK/NRL TPM
High CHUK/NRL TPM
Months
d
Overall Survival
Percent survival
Low HIF-P4H-1 Group
High HIF-P4H-1Group
Months
e
HIF-P4H-1
IKKα
Dapi
Merge
Low Aggressive
High Aggressive
IKKα/HIF-P4H-1 (Ratio)
ns
*
HIF-P4H-1
IKKα
20 µm
f
scrm
siHIF1α
kDa
IKKα
HIF1α
α-Tubulin
85
93
50
Relative IKKα levels
ns
scrm
siHIF1α

#### Supplementary Figure 2

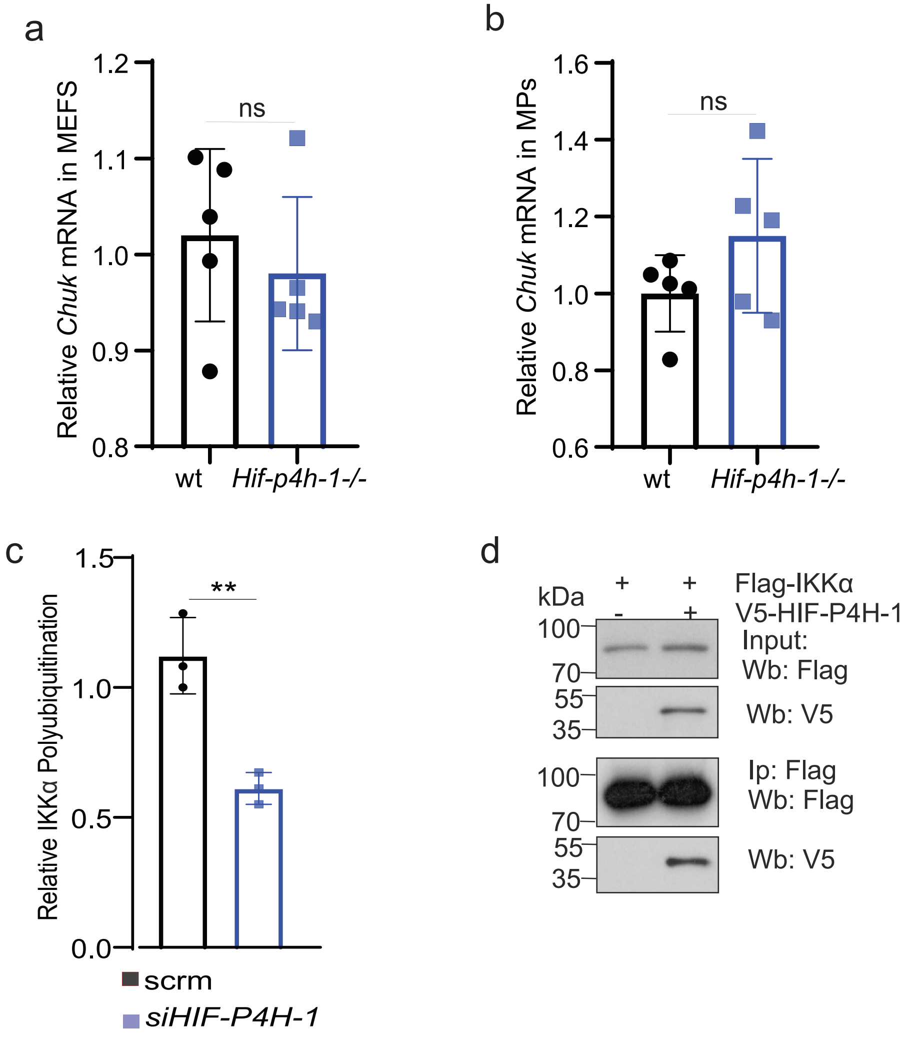

a

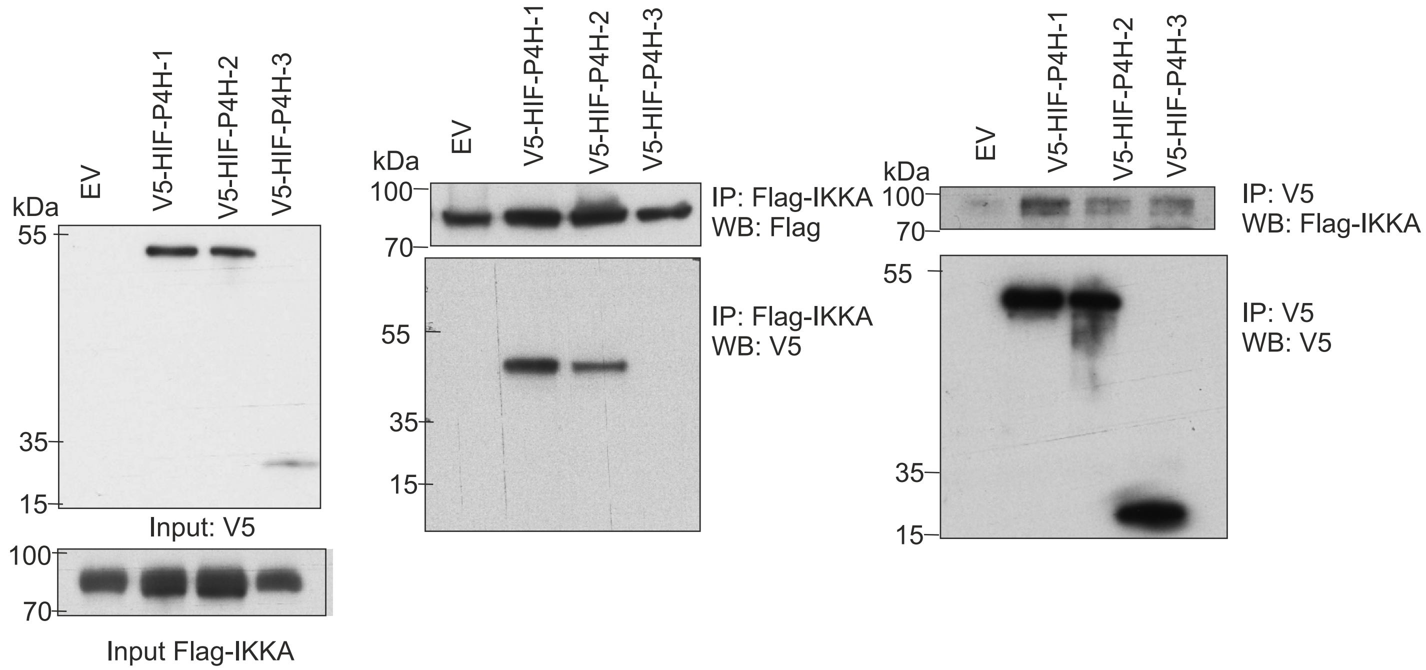


b

Human IKKα : 180 S F V G T **L Q Y L A P** E L F E 194

Mouse IKKα: 180 S F V G T **L Q Y L A P** E L F E 194

The canonical HIF-P4H consensus motif **LXXLAP** corresponds to **LQYLAP** at residues **185–190** in both species

Human IKKα: 366 K **P** A S Q C V L D G V R G C D 380

Mouse IKKα: 366 K **P** A S Q C V L D G V R G C D 380

**P**367 (validated hydroxylation site by LC-MS)

c

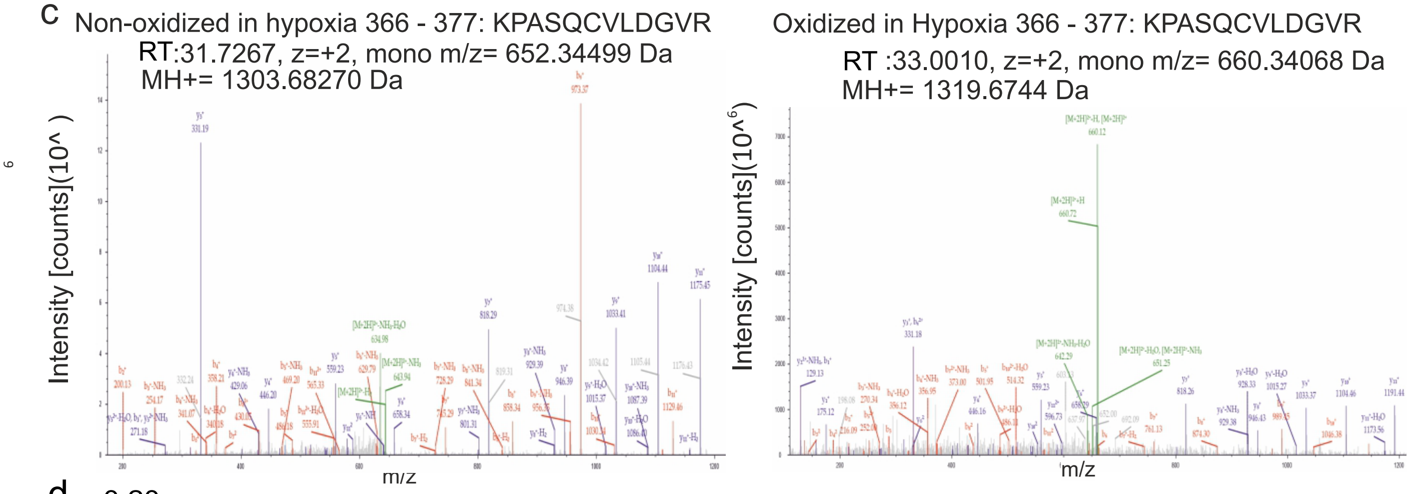


d

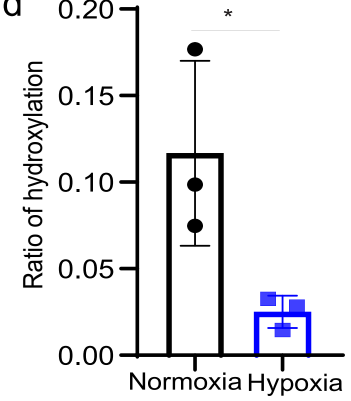

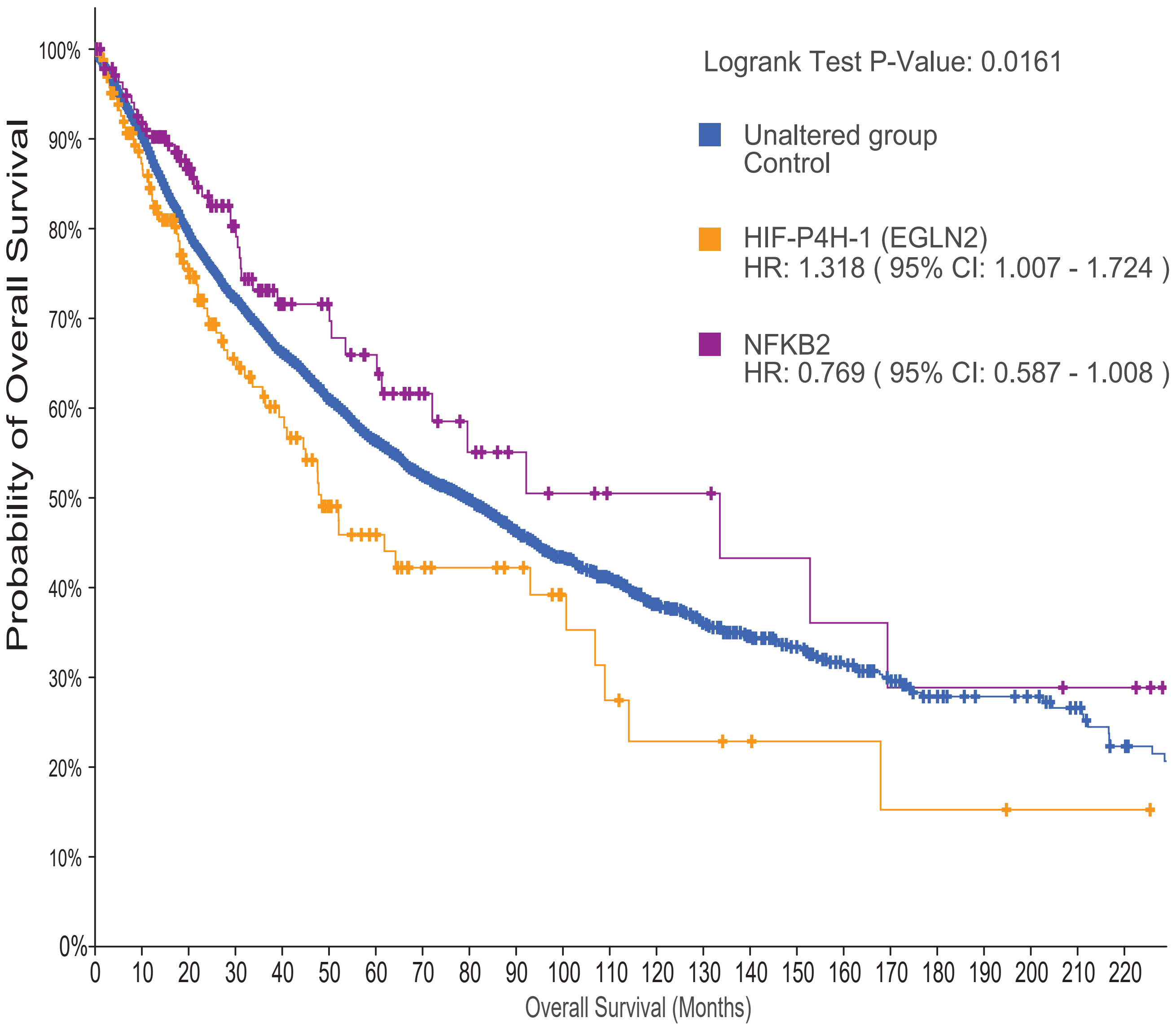


Number at risk (n)

| | | | | | | | | | | | | |
|---|---|---|---|---|---|---|---|---|---|---|---|---|
| Unaltered group | 10378 | 5859 | 2972 | 1680 | 953 | 530 | 294 | 161 | 98 | 59 | 46 | 30 |
| EGLN2 | 171 | 91 | 51 | 26 | 17 | 10 | 5 | 4 | 3 | 2 | 1 | 1 |
| NFKB2 | 140 | 89 | 44 | 31 | 16 | 10 | 8 | 6 | 5 | 4 | 4 | 3 |

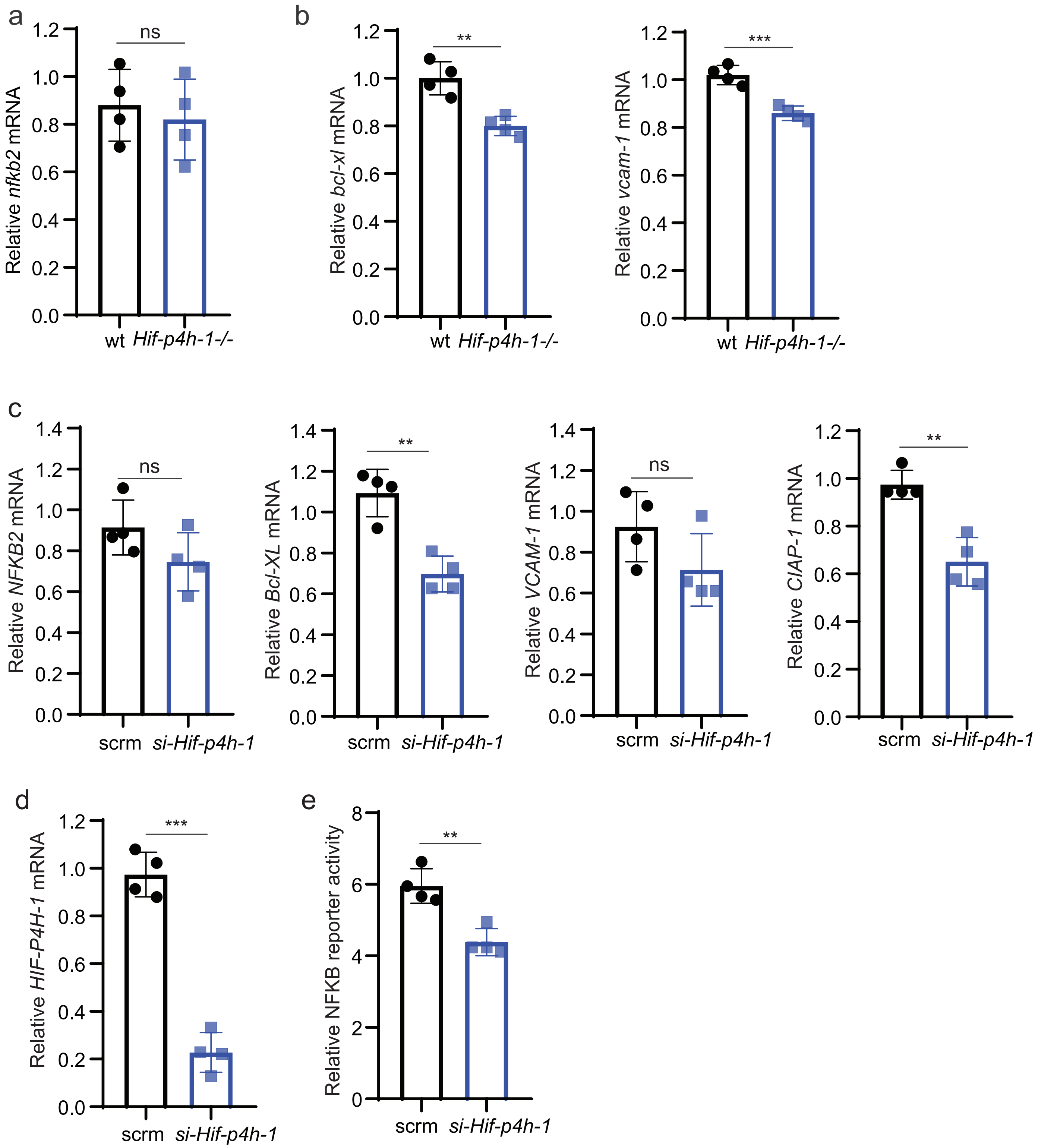
a
b
c
d
e
ns
**
***
Relative nfkb2 mRNA
Relative bcl-xl mRNA
Relative vcam-1 mRNA
wt Hif-p4h-1-/-
Relative NFKB2 mRNA
Relative Bcl-XL mRNA
Relative VCAM-1 mRNA
Relative CIAP-1 mRNA
scrm si-Hif-p4h-1
Relative HIF-P4H-1 mRNA
Relative NFKB reporter activity

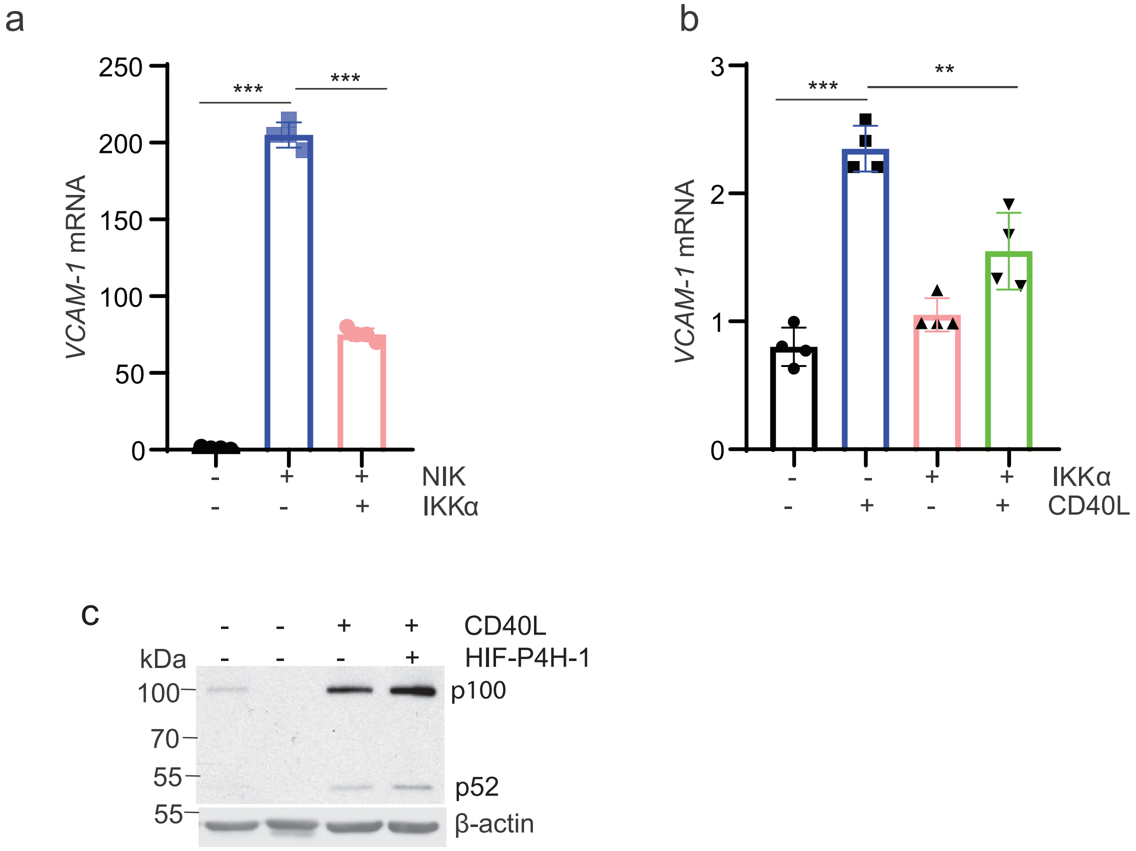
a
VCAM-1 mRNA
250
200
150
100
50
0
***
***
NIK
IKKα
b
VCAM-1 mRNA
3
2
1
0
***
**
IKKα
CD40L
c
CD40L
HIF-P4H-1
kDa
100
70
55
55
p100
p52
β-actin

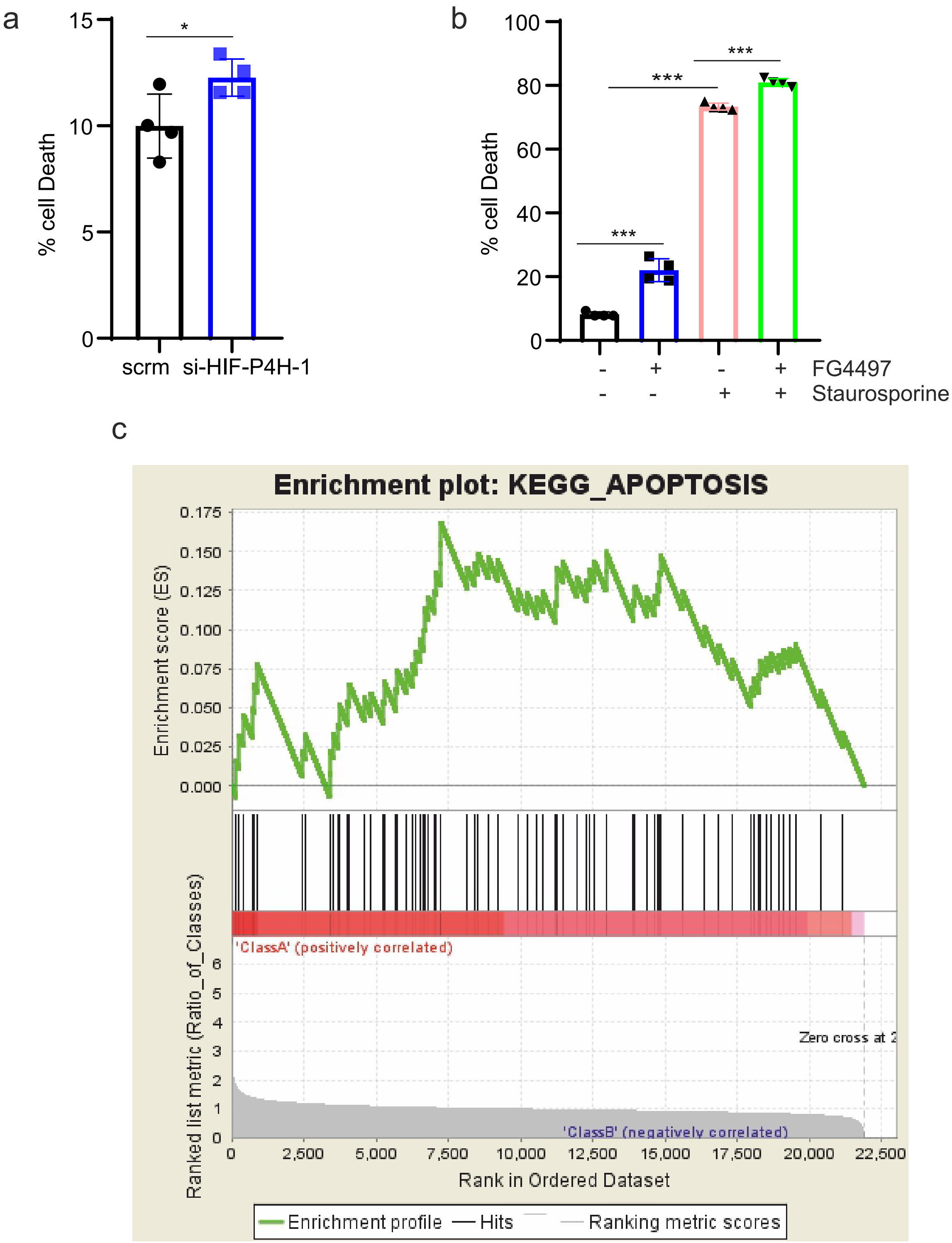
a
% cell Death
0
5
10
15
*
scrm
si-HIF-P4H-1
b
% cell Death
0
20
40
60
80
100
***
***
***
- + - + FG4497
- - + + Staurosporine
c
Enrichment plot: KEGG_APOPTOSIS
Enrichment score (ES)
0.000
0.025
0.050
0.075
0.100
0.125
0.150
0.175
Ranked list metric (Ratio_of_Classes)
'ClassA' (positively correlated)
Zero cross at 2
'ClassB' (negatively correlated)
0
1
2
3
4
5
6
0
2,500
5,000
7,500
10,000
12,500
15,000
17,500
20,000
22,500
Rank in Ordered Dataset
Enrichment profile
Hits
Ranking metric scores

### Supplementary Figure 8

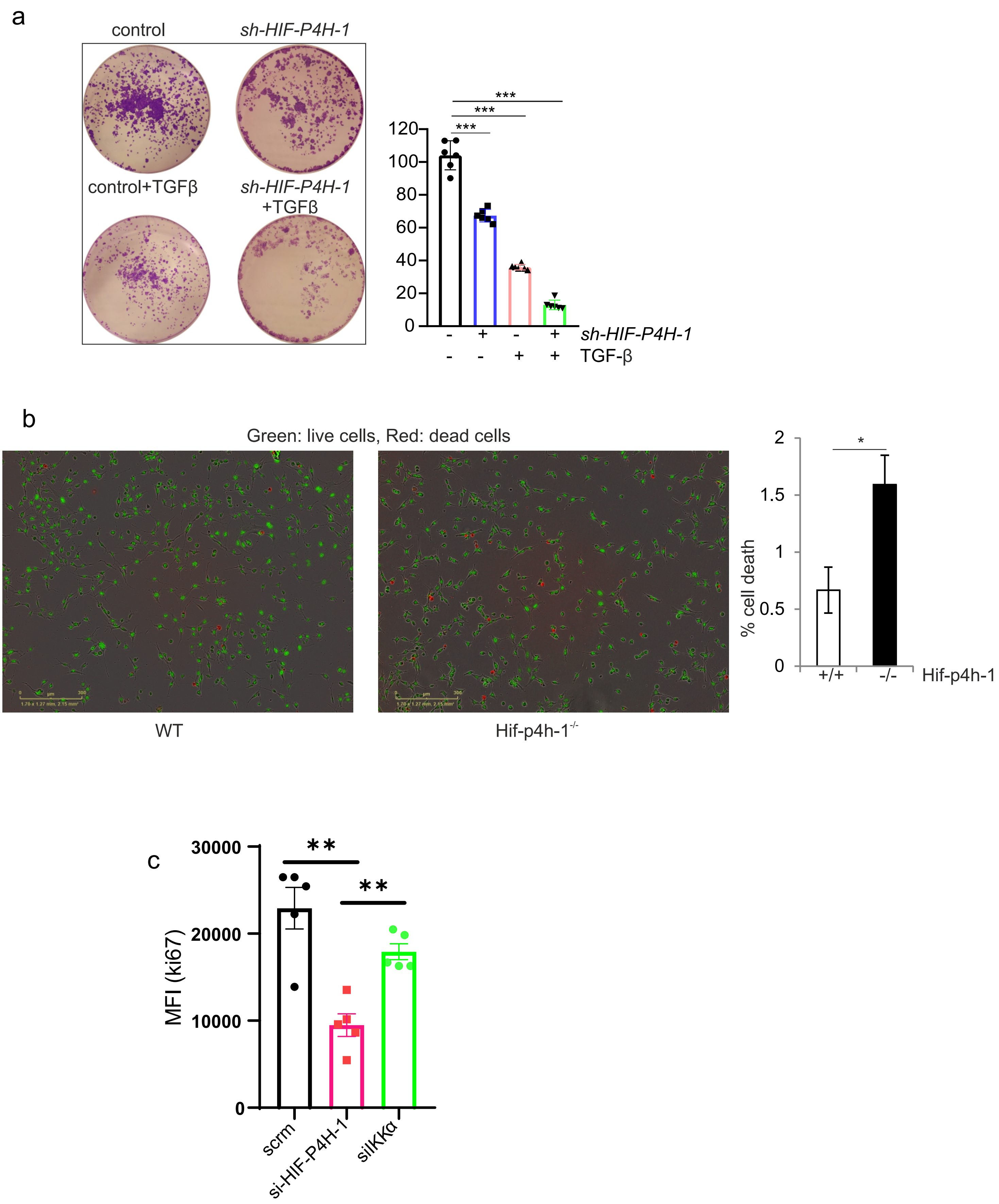
